# Towards a complete phage tail fiber structure atlas

**DOI:** 10.1101/2024.10.28.620165

**Authors:** Victor Klein-Sousa, Aritz Roa-Eguiara, Claudia S. Kielkopf, Nicholas Sofos, Nicholas M. I. Taylor

**Affiliations:** Structural Biology of Molecular Machines Group, Protein Structure & Function Program, Novo Nordisk Foundation Center for Protein Research, Faculty of Health and Medical Sciences, University of Copenhagen, Blegdamsvej 3B, 2200 Copenhagen, Denmark; Core Facility of Integrated Microscopy at University of Copenhagen (CFIM), Copenhagen, Denmark

**Keywords:** Tail Fiber, Phage, Structure Prediction, Domain identification, Cryo-EM

## Abstract

Bacteriophages use receptor-binding proteins (RBPs) to adhere to bacterial hosts. Understanding the structure of these RBPs can provide insights into their target interactions. Tail fibers, a prominent type of RBP, are typically elongated, flexible, and trimeric proteins, making it challenging to obtain high-resolution experimental data of their full-length structures. Recent advancements in deep learning-based protein structure prediction, such as AlphaFold2-multimer (AF2M) and ESMfold, allow for the generation of high-confidence predicted models of complete tail fibers. In this paper, we introduce RBPseg, a method that combines monomeric ESMfold predictions with a novel sigmoid distance pair (sDp) protein segmentation technique. This method segments the tail fiber sequences into smaller fractions, preserving domain boundaries. These segments are then predicted in parallel using AF2M and assembled into a full fiber model. We demonstrate that RBPseg significantly improves AF2M v2.3.2 in terms of model confidence, running time, and memory usage. To validate our approach, we used single-particle cryo-electron microscopy to analyze five tail fibers from three phages of the BASEL collection. Additionally, we conducted a structural classification of 67 fibers and their domains, which identified 16 well-defined tail fiber classes and 89 domains. Our findings suggest the existence of modular fibers as well as fibers with different sequences and shared structure, indicating possible sequence convergence, divergence, and domain swapping. We further demonstrate that these structural classes account for at least 24% of the known tail fiber universe.

## I. Introduction

Bacteriophages are the best-characterized viruses targeting microbes, however, our structural understanding of their receptor-binding proteins (RBPs) remains limited^1,2^. RBPs play a crucial role in identification and attachment to host receptors, including outer membrane proteins (Omp), exopolysaccharides and flagella^3–5^. RBPs are frequently classified into two major groups, tail fibers and spikes. In matured virions, these proteins usually form long homo-trimeric complexes, and attach N-terminally to the tail, most often the baseplate, neck or to other fiber-like proteins and adaptors. Tail spikes and tail fibers differ mostly on their modes of action. Conventionally, RBPs are classified as spikes if they present a depolymerase activity^6–8^.

RBPs are known to be modular, and have domains that are hypothesized to be exchangeable between different phage species through horizontal gene transfer^9–11^. Often, RBPs have an N-terminal part that is sequence and structurally conserved and that binds to the phage baseplate^12–14^, and a C-terminal tip that functions either to identify and adhere to the host receptor^15^, or that connects to other RBPs, forming a complex host receptor detection apparatus^16^. For instance, phage T4 has two tail fiber sets: a short tail fiber (STF) formed by gp12, responsible for pseudo-irreversible binding to a secondary receptor (lipopolysaccharides, LPS), and a long tail fiber (LTF) formed by gp34, gp35, gp36 and gp37, which adheres to Omp^17^. *Siphoviridae* phages, such as T5 and lambda^18,19^, and some *Podoviridae* phages^20^, exhibit lateral tail fibers, and a central tail fiber that is connected to the baseplate hub protein.

Recent biotechnological studies demonstrated the possibility of creating chimeric RBPs to target novel hosts by segmenting the fibers in precise hotspots^21,22^, keeping their oligomeric state stable. As RBPs are responsible for specific interactions with the hosts, a broad understanding of RBP types and functions plays a critical role in medical applications of phages, such as engineered phage therapy^23^. Insight into their structural organization advances this knowledge.

Currently, the experimental methods modelling complexes face limitations, especially when working with elongated and flexible RBPs. Obtaining high-resolution maps using single-phage cryo-electron microscopy is challenging due to conformational variability. For similar reasons, obtaining a good quality crystal of a full fiber is an obstacle for structural elucidation by X-ray crystallography. Nevertheless, recent advancements in computational protein structure prediction, such as AlphaFold^24^, ESMfold^25^ and AlphaFold-multimer (AF2M)^26^ have made it feasible to generate reliable models for proteins and protein complexes.

Modeling large protein complexes using standard AF2M pipelines still faces computational limitations, mostly due to hardware and memory issues, resulting in models with large low-confidence regions, clashes, and long computing times. Strategies to address the challenge of building large complex models using AF2 were previously proposed^27,28^. But current methods were not specifically designed for elongated fiber-like structures.

Previous studies have used AF2M to predict the structure of RBPs^9,29^, where human expert information was used to manually segment the sequences into smaller pieces, model the pieces individually, and assemble the fractions into a final model. However, an automated and systematic way of determining the location and number of segmentation points is lacking.

Here, we propose a novel pipeline, RBPseg, that defines major domains and segmentation hotspots based on tail fiber primary structure using ESMfold and a novel sigmoid distance pair function. RBPseg then predicts tail fiber fractions in parallel using AF2M and assembles full fiber models. This systematic approach overcomes computational limitations, improves performance, and yields more reliable models for large tail fibers. Overall, our approach provides a robust solution for predicting, classifying, and characterizing phage tail fibers, offering insights into their structural properties and potential functional roles.

We implemented RBPseg for a selected range of model phages and classified the predicted RBPs based on structural similarity and domain specificity. These models represent 24% of the known tail fiber universe and provide as valuable information for analyzing the phage toolbox for host attachment by identifying structurally related domains.

## II. Methods

### II.1 Sigmoid distance pair function (sDp)

The sigmoid distance pair function (sDp) is used to segment a protein structure model into smaller fractions. Given a pair of residues in the model (A and B, where A ≠ B), we define the sDp function as follows:

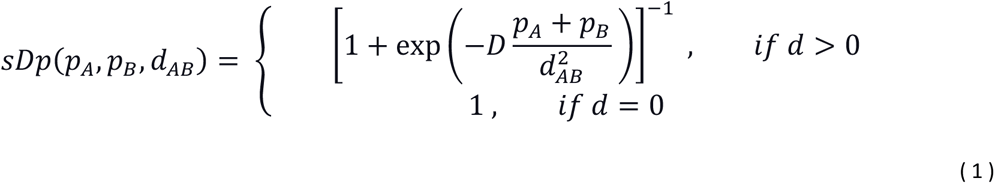

Where *p_A_* and *p_B_* are the normalized pLDDT of residues A and B from the ESMfold or AF model, ranging from 0 to 1, and d is the C_α_-distance of the residue pair, multiplied by the pair distance constant (D) assumed to be 1 Å^-2^. The sDp is a continuous and differentiable function for d > 0. For computational purposes, if d is equal to zero, we define the sDp as equal to one. Hence, this function is restricted to the interval of (0.5,1).

### II.2 Fraction module and pseudo-domain identification

In a protein model with predicted per-residue LDDT, two residues will be considered part of the same pseudo-domain if they are in proximity (low *d_AB_*) and share similar pLDDT values. This assumption allows us to use the sDp to identify possible domains in a protein prediction model, independent of the Predicted Aligned Error (PAE) matrix.

Given a protein structure model containing the pLDDT values in the B-factor column, we calculate an all-against-all sDp matrix. This matrix undergoes dimensionality reduction using UMAP^30^, creating a 2D-latice representation of the 3D structure. Clustering methods (k-means or HDBSCAN^31^, the latter was used for this study) are used to identify neighboring residues and pseudo-domain interfaces. The number of clusters is self-optimized by calculating the silhouette score^32^ for different k (by default with a maximum k of 10).

### II.3 Sequence segmentation

After defining the fraction borders, two consecutive domain sequences are stored in a fraction file, ensuring at least one domain overlap in two consecutive FASTA files. A minimum cut-off of 50 residues per monomeric sequence is required for each FASTA file. If a domain has fewer than 50 residues, the subsequent domain is appended to the file until the sequence reaches a minimum of 50 residues. This precaution is taken to mitigate the risk of low confidence predictions associated with very short peptides.

### II.4 ESMfold and AlphaFold2-multimer models

The monomeric models for the fibers are generated using a standard installation of ESMfold-v1, with maximum token per batch of 1024, 4 recycles and chunk-size of 32. AlphaFold-multimer v2.3.1 (AF2M) is used to predict models of protein fractions with full MSA search, and three recycling cycles without a relaxation step. Same conditions were applied for the models of whole fibers (without segmentation). All predictions were run on an NVIDIA HGX™ A100.

### II.5 Assembling of tail fiber models

Two consecutive fiber fractions models are superimposed based on their overlapping region. The residue pair with the minimal per-residue RMSD is selected as the joint point. The inter-model chain pair is assigned by minimizing the peptide bond length. To prevent poor assignments due to inadequate superposition, an additional restriction imposes that the new peptide bonds should not cross into the inner space between chains, which is defined by a sphere centered inside the triangle formed by the trimer of the next C-alpha residues in the sequence. The sphere radius is equivalent to 1/3 of the triangle height. This process is done for all generated fraction models, and the pair with optimal RMSD is selected for merging. Subsequently, the merged model undergoes Amber relaxation.

### II.6 Selection of BASEL Collection Tail fibers

The genomes of all phages in Maffei et al.^33^ were accessed through their GenBank ID, and their proteome was collected at UniProtKB^34^. Fibers were selected based on the consensus of PhaNNs^35^ and PhageRBPdetect^36^. A threshold of 8 was used to select positive predictions of PhaNNs. The selected sequences were clustered hierarchically based on sequence identity retrieved from ClustalΩ^37,38^ (default parameters). 67 sequences were randomly selected, respecting the proportions of each major group, and relabeled as RBP_{index}.

### II.7 Structural and sequence-based comparison and clustering of tail fibers

The RBPseg models for all analyzed fibers were clustered based on their pair inter-model TM scores^39^. US-align^40^ (fast-mode) was employed to run all-against-all protein pairs. Utilizing the TM-score, we generated a quasi-symmetric squared structural identity matrix. On this matrix, clusters were calculated using the spectral clustering method. The optimal number of clusters was estimated to maximize the Silhouette score (Sil) and the minimal inner cluster TM-score (icTM), and to minimize the maximum cluster size (CS) and the standard deviation (std) of icTM. This was done by calculating the normalized structural clustering similarity metric (SM) for each cluster number (n) as follow:

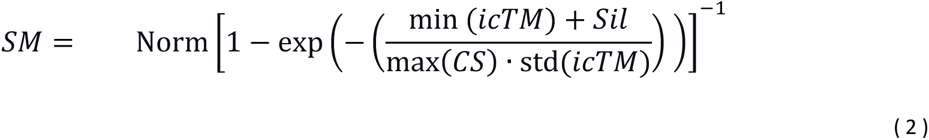

We model the normalized predicted SM (pSM) given n as an exponential decay function:

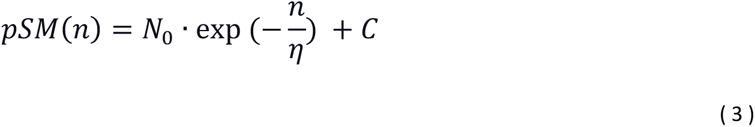

The second order derivative of pSM provides the deacceleration decay, which converges to zero in the limit *n* → ∞. The optimal n was selected for each case imposing: *pSM*′′(**n*_op_*) = *ε*, where *ε* << 1.

The phylogenetic tree was constructed using a ClustalΩ alignment and implementing ETE Toolkit v3.1.3^41^. The sequence classes were generated by applying MMseqs2^42^ with a minimal sequence identity of 0.4 and coverage of 0.5.

### II.8 Fiber domain search and classification

The models for the fibers were divided into domains by applying the sDp approach. Residues that were not classified in a cluster were automatically appointed to the cluster of the nearest residue. The individual domains underwent structural comparisons, first all-against-all, with the same approach as described in II.7, subsequently all-against RCSB-PDB^43^ (RCSB.org date: 2023-01-12) by implementing Foldseek^44^. The full fiber sequences were analyzed on InterPro against all databases ^45,46,47^.

### II.9 Comparison of structural classes with known tail fibers and tail fiber atlas

Sequences of annotated fibers were found on UniProtKB^34^. We filtered the results containing ("tail fiber" or "tail fibre") AND (length: [300 TO 3000]) AND (organism_name:phage) NOT chaperone. An MSA and an HMM profile was created for each structural fiber class, excluding TC5 and TC17. Hmmsearch was implemented to search for sequence homologs against the UniProt selected sequences. The tail fiber atlas was created by implementing ipysigma-v0.24.0. ^50^.

### II.10 Statistical analysis

To test the effect of segmentation of the fibers on the precision of the AF2M predictions, we used a Mann–Whitney *U* test on the per-residue pLDDT distributions of RBPseg results and AF2M. The null hypothesis is that for randomly selected values of the pLDDT populations, the probability of pLDDT_RBPseg_ being greater than pLDDT_AF2M_ is the same as for the opposite (pLDDT_RBPseg_ lower than pLDDT_AF2M_). We further compared the mean total pLDDT per model using a t-test for the means of the two independent datasets, with the null hypothesis being that they were equal. Both tests were mono-caudal, and the null hypothesis was rejected for p-value < 0.01. The calculations were made by implementing SciPy.stats v1.11.4^51^, and p-values were calculated with an asymptotic approximation. The same tests and conditions were applied for the benchmark metrics (running time, MaxRSS, mean MSA coverage).

### II.11 Phage purification

The BASEL phages were propagated and purified using standard phage purification protocols^33^. *E. coli* MG1655 ΔRM was used as a host. A primary stock of the phage was prepared using a double layer agar method (soft layer: 0.6% LB agar; hard layer: 1% LB agar). A small batch lysate (50 mL) was prepared by growing the host until OD_600_ of 0.2 in LB media supplemented with 20 mM MgSO_4_ and 5 mM CaCl_2_. The primary stock of the phage was incubated with the host in a MOI < 1 for 4-5 hours until OD_600_ dropped below 0.1. Next, 1:100 (v:v) chloroform was added for 15 minutes. The supernatant was used to infect 1 L of host in similar conditions as before. After lysis, 5 μg/mL of DNase 1 and RNase A were added for 1 hour, followed by incubation with 30 g/L NaCl and 75 g/L PEG 8000 overnight at 4 °C. The precipitated phage was pelleted at 15,000 x g for 60 minutes and resuspended in 5 mL of SM buffer (100 mM NaCl, 8mM MgCl, 50 mM Tris-HCl pH 7.5). Chloroform was added at a ratio of 1:1 (v:v), and the sample was inverted until it became homogeneous and centrifuged at 6,000 x g for 15 minutes. The supernatant was loaded onto an OptiPrep™(Sigma-Aldrich) gradient (50%-10%) and was centrifuged at 150,000 x G for 18 h. Fractions of each gradient containing the phage (between 0.5 to 1 mL) were diluted to 5 mL and were ultracentrifuged 72,000 x G for 1 hours. The phage pellet was resuspended in 100 μL of SM buffer. The phages were further purified and concentrated by pelleting at 20,000 x G for 45 minutes and resuspension in 50 μL of SM buffer.

### II.13 Cryo-electron microscopy data collection

Phage samples were applied to R2/2 grids (Bas49 and Bas54 on Quantifoil and Bas36 on UltrAuFoil grids) with 2 nm continuous carbon layer and vitrified in liquid ethane cooled by liquid nitrogen, using a Vitrobot Mark IV robot. Individual datasets were collected on a Thermo Scientific Krios G2 with a Falcon 4i Direct Electron Detector and Selectris X Imaging filter at a dose of 40 e/A^2^ and a pixel size of 1.2 Å.

### II.14 Cryo-electron microscopy data processing

All cryo-EM data was processed using CryoSPARC v4.5.1^52^ and ChimeraX v1.6.1^53^. All phages datasets were preprocessed similarly: Movies were patch motion corrected (default parameters) and the contrast transfer function (CTF) was estimated (patch CTF estimation with default parameters), followed by discarding micrographs with worse resolution than 15 Å and outliers on ice thickness, yielding 5663, 6591, and 5142 movies for Bas36, 54, and 49 respectively. Baseplates were picked using the blob picker tool. Particles were extracted with a box size of 750 px and Fourier cropped to 300 px. 2D classification was applied to obtain projections of the baseplate, that was further used for template picking. New picks were extracted with box size of 750 px and Fourier cropped to 300 px. 2D classification was used to discard “junk” classes, followed by ab-initio and heterogeneous refinement to select only baseplates. Homogenous refinements were used to obtain models of phage baseplates and masks were created around on of the fiber sets for each phage. Subsequent iterations of local refinements, 3D classification, recentering, and CTF refinements around the regions of interest led to the final maps of the Bas36 LTF proximal region and fiber networks of Bas49 and 54.

Model to map fit was evaluated using comprehensive cryo-EM validation in PHENIX^54^ (v 1.21), using a resolution cut-off (local resolution in the fiber area) of 9.5 Å.

## III. Results and discussion

### III.1 RBPseg workflow

RBPseg is an implementation of two protein structure prediction approaches, ESMfold and AF2M, to handle large fiber-like multimeric structures (**Fig. 1a, Supplementary Fig. 1**). Specifically, RBPseg was designed to improve predictability of full quaternary structure of phage tail fibers and spikes. Overall, RBPseg uses tertiary structural information (pseudo-domains), obtained from an ESMfold model, to automatically fragment protein sequences while preserving key features of the fiber. The fractions are defined by calculating the sigmoid distance pair (sDp) matrix and performing unsupervised clustering. FASTA files are created containing two or more consecutive fractions, with a one pseudo-domain overlap in two sequential files. The models for the protein fractions are predicted using AF2M, without relaxation and with three recycling steps. Models for all fractions are superimposed based on their overlaps, and the pairs with the smallest RMSD are sequentially merged into a full fiber model, respecting the correct chain connectivity, and the final model is amber relaxed^55,56^.

**Fig. 1.**
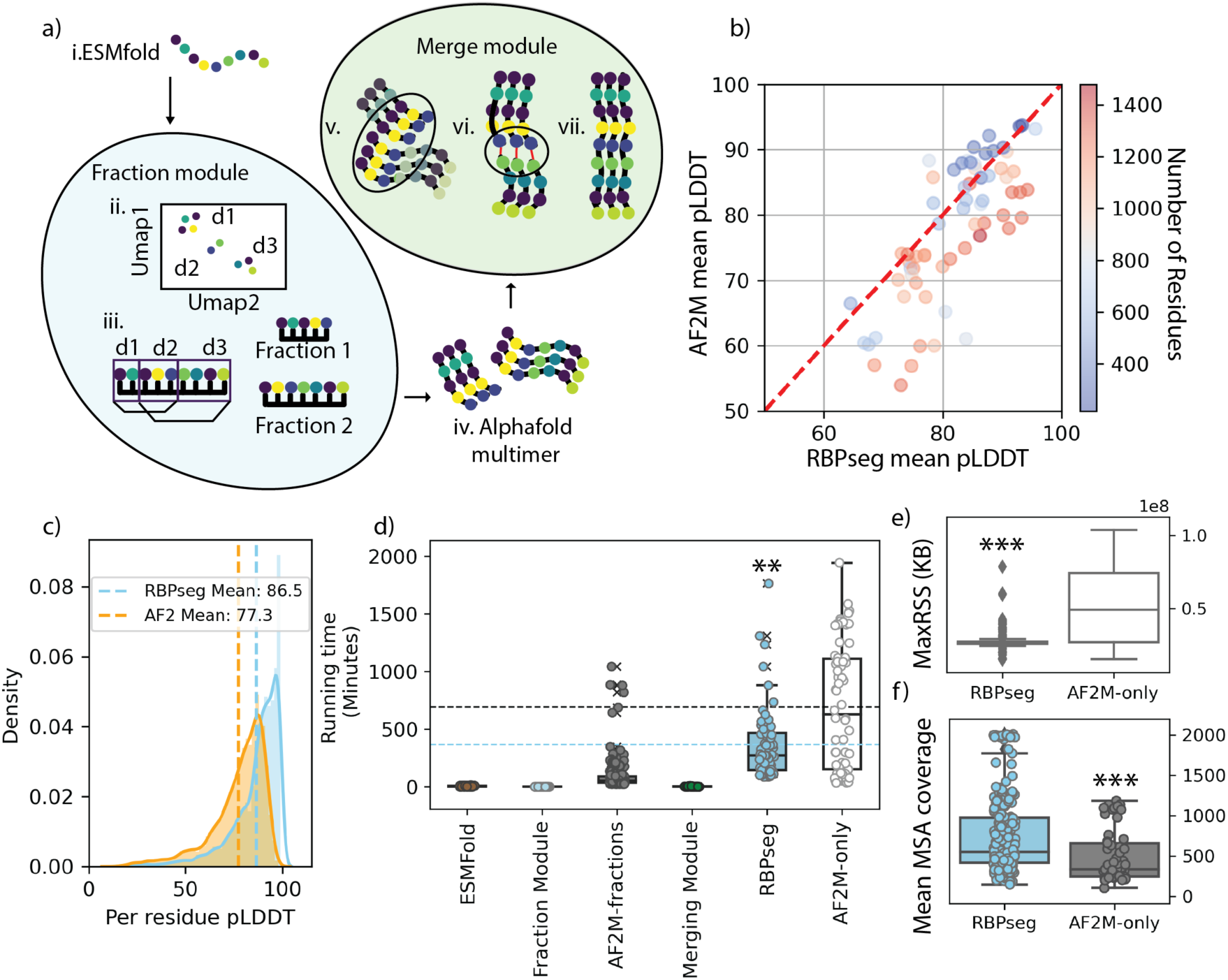
– RBPseg workflow increases confidence of AlphaFold2 models. a) The RBPseg pipeline starts by inputting an ESMfold prediction (i) of the monomeric RBP. The Fraction module first applies the sDp approach to find possible domains in the structure (ii) and the sequence of these domains are arranged in consecutive pairs to create fractions (iii). The fraction sequences are modeled as trimers by AF2M (iv.). These resulting modules are input into the Merge module, that (v.) superimposes the overlap domains, (vi.) pairs and connects the chains, and (vii.) runs an amber relaxation. (b-e) Benchmarking of RBPseg against regular AlphaFold-multimer v2.3.1. b) Per-model mean pLDDT comparison between AF2M and RBPseg, with individual dots colored by sequence length. c) The density distribution of per-residue pLDDT of RBPseg and AF2M. d) Running time (minutes) of each component in the RBPseg pipeline. From left to right: Monomeric-ESMFold prediction, sDp module, AF2M prediction of single fraction (AF2M-seg), merging and relax module, RBPseg, and running time of AF2M prediction of the full fiber sequence (AF2M-only). e) Maximum RSS (Resident Set Size) of RBPseg against AF2M. f) Per-residue mean MSA coverage of RBPseg fractions compared to AF2M, showing that the MSA coverage is significantly larger for RBPseg fractions. Statistical significance is indicated by a one-tailed Mann–Whitney U test (**p < 0.01, ***p < 0.001).

### III.2 RBPseg improves AlphaFold2-multimer v2.3 for prediction of tail fibers

We benchmarked the RBPseg pipeline and AF2M-v2.3 using selected tail fibers of well-studied phages^33^. Initially, 432 sequences were identified as RBPs and tail fibers (**Supplementary Fig 2a**), from which we performed a sequence-based hierarchical clustering, that subdivided the fibers into four major groups (**Supplementary Fig 2b-d**). We then randomly selected representatives from each group, respecting their relative abundance, totaling 67 sequences (**Supplementary Fig 2e**). We ran RBPseg and AF2M-v2.3 for all sequences.

The performance comparison demonstrates that the RBPseg workflow improves AF2M in terms of model confidence, running time and memory usage (**Fig. 1b-f**). The models generated with RBPseg show higher mean-pLDDT scores, especially for longer sequences, and significantly higher mean per-residue pLDDT (**Fig. 1b and c**). Moreover, we verify a significant decrease in total running time (**Fig. 1d**). A full tail fiber model was acquired in 6.24 hours on average, using RBPseg workflow, whereas AF2M had a mean running time of 11.56 hours, with a significantly higher distribution (Mann-Whitney U test, p-value of 0.2×10^-15^). Furthermore, the modelling of fractions by RBPseg used significantly lower peaks of memory usage (maximum Resident Set Size) than AF2M (**Fig. 1e**). These results suggest that AF2M alone is more memory-intensive than RBPseg, which tries to improve computational efficiency by segmenting the sequences. The shorter sequences in the fractions also led to improved per-residue MSA coverage (**Fig. 1f**), indicating a better signal-to-noise ratio for the overall MSA, which is reflected in the higher accuracy of RBPseg models.

To further validate the RBPseg models, bacteriophages *Escherichia phage* Paracelsus (Bas36), *Escherichia phage* MaxTheCat (Bas54) and *Escherichia phage* EmilHeitz (Bas49) were purified in high titer and cryo-EM datasets were collected on them. After local refinements, we obtained low-resolution (7.3∼9.5 Å, **Supplementary Table 1**) maps from five distinct tail fibers (**Fig. 3**). We modelled and fitted the predicted models into the maps and obtained high cross-correlation for all of them (**Supplementary Fig. 4a-b**). Notably, the real curvature of the fibers could not be modelled, which resulted in badly fitted regions and low cross-correlation. The cross-correlation increased when the pseudo-domains of the models were fitted, demonstrating that the RBPseg models preserved the domain organization.

### III.2 The Sigmoid distance pair function is a general method to identify protein pseudo-domains given a protein prediction model

To test the general applicability of the sDp approach for identifying protein regions, we used a dataset of a thousand randomly selected protein models from the AlphaFold database^57^ for *E. coli* (**Supplementary Fig. 3**). We confirmed that the all-against-all residue pair sDp matrix exhibits a strong negative Pearson correlation with the predicted aligned error (PAE). We calculated the pair distance constant (D) that maximizes the correlation between sDp and PAE by fitting a 4th order polynomial, as 9.87 Å^2^. At this value, we estimated a correlation of −0.71 (**Supplementary Fig. 3c-d)**. The sDp-PAE correlation is dependent on sequence length and is reduced for structures with low mean PAE (**Supplementary Fig. 3f-g**), as sDp is a metric that preserves the spatial distribution of pairs independent of PAE.

We also evaluated the impact of different sDp calculation methods. In the first modification, the exponential term was divided by the absolute Euclidean distance of the residue pairs instead of the square distance. In the second modification, we used the product of the pair pLDDT values instead of the sum. In both cases, a strong correlation with PAE persisted when 0.1 < D < 1 Å (or Å^2^). The most significant change was observed when performing dimensionality reduction (UMAP) on the resulting matrices. For all sDp models, we observed a 2D lattice projection of the 3D model that preserved the overall structural organization, which was not found for the PAE (**Supplementary Fig. 3**). For the purposes of this study, no significant differences were observed when applying the different sDp variants, although they might result in slightly different pseudo-domain organization.

### III.3 The BASEL phages have sixteen well-defined tail fiber structural classes, with shared modules

We further analyzed the dataset that was used to benchmark the RBPseg method by investigating the sequence and structural similarities between the different fibers. At sequence level, we identified 24 unique clusters using mmseq2, with 8 singletons (**Supplementary Table 2**).

To assess structural similarity, we performed a TM alignment of all-versus-all tail fiber models (**Fig. 2, Supplementary Fig. 5-6**), removing four outliers: two poorly aligned models and two sequence singletons likely misidentified as fibers. The resulting TM matrix was spectrally clustered into 18 classes (TC0-TC17). The optimal number of clusters was determined using the predicted similarity metric (pSM, **Supplementary Fig. 7**. To distinguish true classes from randomly assigned classes, we calculated the mean TM score (<TM>, **Fig. 2b**) for all TC and tested if they were greater than a randomly assigned class average (single tail t-test, p-value < 0.01). This resulted in the exclusion of two classes (TC5 and TC17; **Supplementary Fig. 6a-b**). The mean inner TM score within the well-defined classes was 0.59, whereas TC5 and TC17 had mean scores of 0.27, and 0.28, respectively (**Fig. 2b**).

**Fig. 2.**
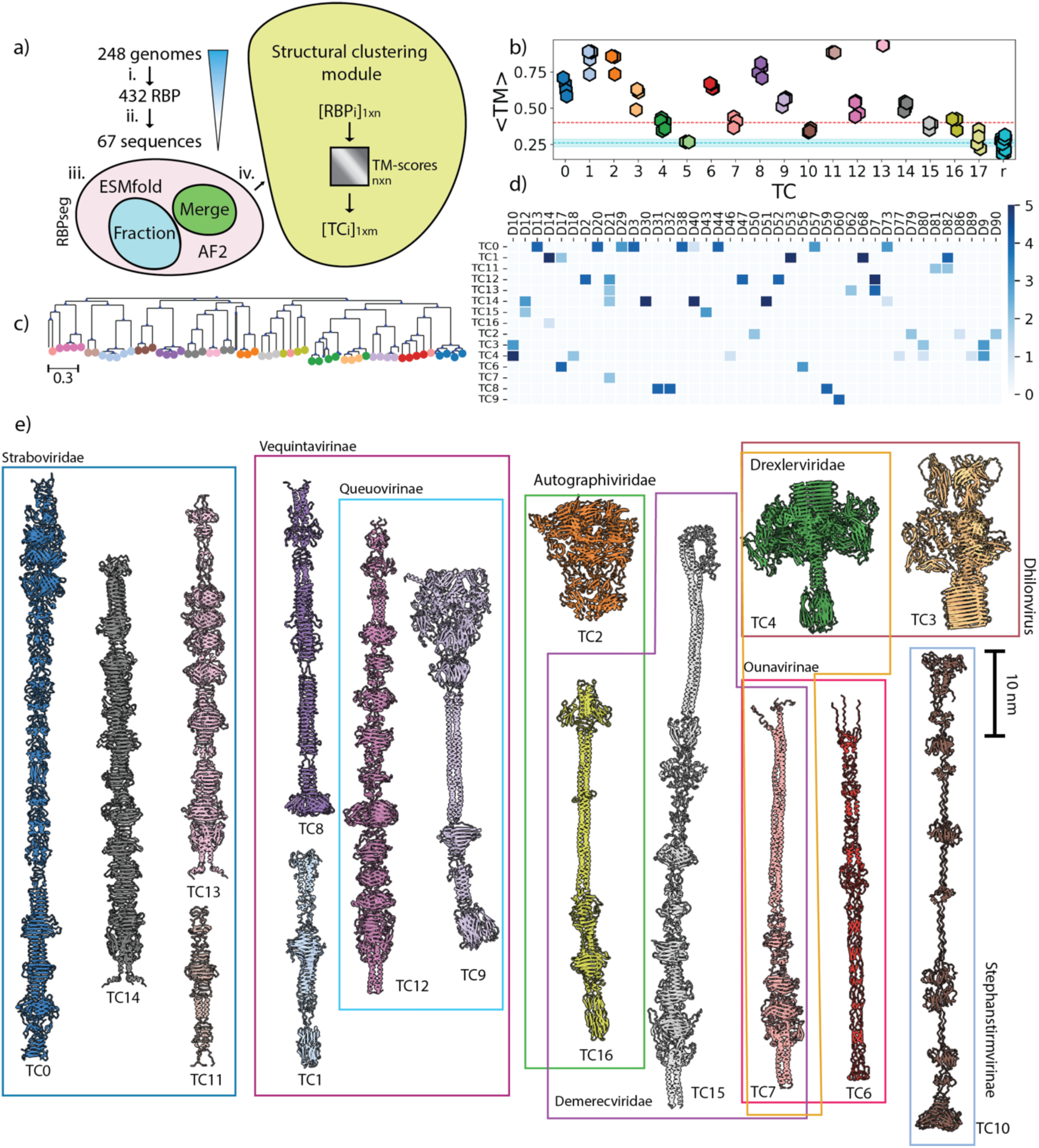
– The BASEL collection RBP classes. a) Summary of the RBP selection process. (i) 432 RBPs were selected among 248 genomes by taking the consensus between PhaNNs and phageRBPdetect. (ii) These RBPs were hierarchically clustered and a total of 67 RBPs were chosen as representatives. (iii) The RBPs structure were predicted using the RBPseg workflow. (iv) The resulting models were classified based on structural similarity. b) The mean TM score value for each TC and a randomly assigned class (r). In cyan, a horizontal line represents the average value of r with an interval of 3 standard deviations of the mean. In red, a horizontal line indicates TM=0.4. c) Unrooted phylogenetic tree based on sequence for the selected 67 RBPs, terminal nodes colored based on the TCs, as in (b). d) A heatmap showing the presence of 41 well-defined pseudo-domains (<TM> > 0.7) in different TC classes. The side bar represents the number of pseudo-domains. e) Representatives for each TC class colored based on (b) and grouped according to their phage family, subfamily or genus.

Interestingly, even though TC5 was excluded based on TM score criteria, we observed some similarities among their members. TC5 consists of two elongated tail fibers (RBP_55 and RBP_22), characterized by large coiled-coil domains, a β-sandwich-rich N-terminal region which is common in central tail fibers, and a β-helix at the C-terminus (**Supplementary Fig. 6a**). Despite these structural similarities, the fibers belong to different viral subfamilies (*Tempevirinae* and *Vequintavirinae*) and sequence subfamilies. TC17 is composed of four poorly grouped fibers: two share structural and sequence similarities with TC0/seq9 (**Fig. 2c**), one is classified as seq14, and one is a singleton (**Supplementary Fig. 6b**).

To better annotate and understand potential functions and distribution of each region within the classes, we applied the sDp approach to all models, generating 702 pseudo-domain structures, including all TC classes. We then spectrally clustered all the pseudo-domains with more than 20, and less than 400 residues. These filtered domains were grouped into 88 structural classes (D-classes, **Supplementary Fig. 8a-c**, **Fig. 2d**). Notably, 52 pseudo-domains were not assigned to any class. A total of 41 D-classes had an inner mean TM score above 0.7 (**Fig. 2d**).

Function annotation for each D-class was performed by running Foldseek and Interpro (**Supplementary Fig. 9; Supplementary Table 3**). We identified a total of 117 unique PDB matches with high confidence (alignment TM score > 0.8) for all TC, and 109 matches when excluding bad classes, including 41 shared matches between different TCs. The number of unique matches increases to 2484 at alignment TM score > 0.6. The InterPro analysis retrieved 32 unique signature accessions with high confidence (e-value > 10e-5). No annotation was found for TC8.

### III.5 Overview of the Sixteen Tail Fiber Scaffolds

We next analyzed the sixteen tail fiber scaffolds to obtain insight into their functional differences. TC0 is a gp34-like^16^ class (**Fig. 2e**, **Fig. 3a**), and it was subdivided into 11 pseudo-domains, each fitting the cryo-EM map for Bas36 proximal LTF. This domain organization is identical to the previously classified regions (P1-5), with region P1 and P2 being subdivided into 4 pseudo-domains each (**Supplementary Fig. 9**). TC0 has three regions (two copies of D19 and D38) that share structural homology of with T4’s short tail fiber.

**Fig. 3.**
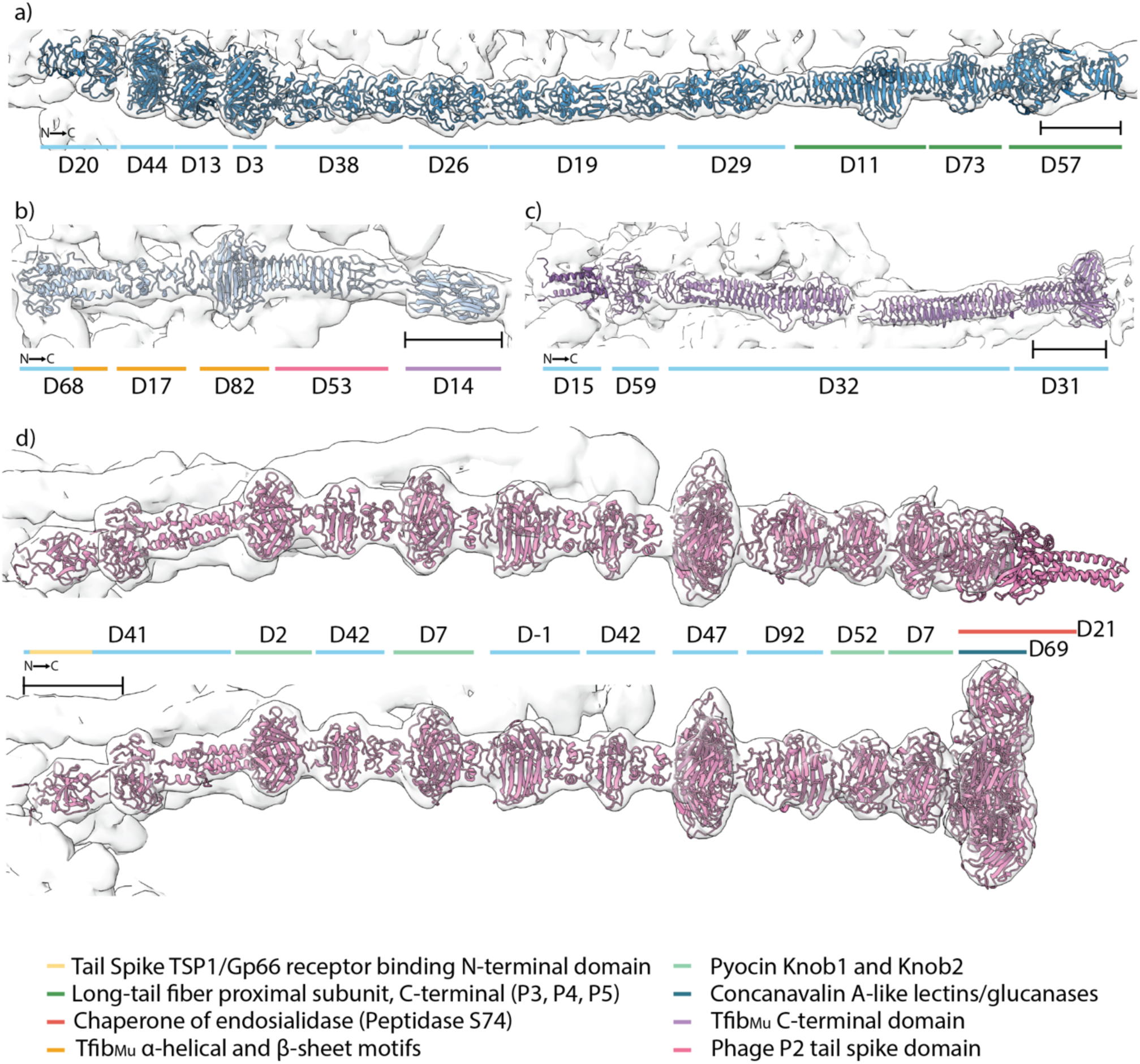
– RBPseg domains fitted into local refined cryo-EM maps. a) Cryo-EM map of matured *Escherichia phage* Paracelsus proximal LTF (RBP39-TC0, EMD-51870) overlapping sDp pseudo-domains for RBP39 model. b,c) Composite cryo-EM maps of *Escherichia phage* MaxTheCat of baseplate and distal fiber region (EMD-51869), with overlapping pseudo-domains of RBP04 (TC1, b) and RBP65 (TC8, c). d) Composite cryo-EM maps of *Escherichia* phage MaxTheCat (top, EMD-51869) and *Escherichia* phage EmilHeitz (bottom, EMD-51868) of baseplate wedge and distal fiber region. Pseudo-domain regions colored based on sequence (Interpro hits with e-value < 10^-5^) or structural homologs (Foldseek hits with alntmscore > 0.6). Skyblue represents pseudo-domains with no siginificant hits.

TC11, TC13, and TC14 are also present in *Straboviridae* phages (**Fig. 2e**). TC11 is a homolog of T4’s gp36 and TC14 is a homolog of gp37. TC13 has a novel fiber architecture, not present in bacteriophage T4. The gp37 homologs in the dataset lack the well-characterized needle domain of phage T4 in their C-termini. Instead, they have a peptidase S74-like domain with a triple–β-helix fold, which aids oligomerization and is known to self-cleave^58^. Such a fold is present in *Salmonella Phage* S16^59^. This C-terminal domain is common across several tail fiber classes (TC7, TC12, TC13, TC14, and TC15) and is found in other phage subfamilies, including *Markadamsvirinae*, *Vequintavirinae*, and *Queuovirinae*.

TC6 comprises fibers with a T4-like needle-shaped receptor-binding tip, all belonging to the *Ounavirinae* subfamily. They are connected to an N-terminal rod by a T4 baseplate protein gp10-like C-terminal domain.

TC1 is a short tail fiber present in *Vequintavirinae* phages. We found a density corresponding to this fiber in both V5 purified phages (**Fig. 3b**). This fiber resembles the phage Mu tail fiber in its C-terminal region, which probably indicates that they require a special tail fiber assembly protein to oligomerize and possibly interacts with exopolysaccharides^60,61^. The same C-terminal homology is also present in a member of TC16 (RBP_50). Interestingly, the D53 region showed Foldseek matches with the P2 central spike gpV^62^, which has a different β-helical region not present in Phage Mu fibers and RBP_50.

TC2 are structural homologs of the T7 gp12 nozzle proteins^13^, which form a hexameric complex, and were misclassified as tail fibers.

TC3, TC4, and TC9 form a larger group of tail needle fibers. The C-terminal region of this type of scaffold is found in phage λ and T5 on their tail tip attachment protein J. The N-terminal fold of these fibers is structurally related to the T5 baseplate hub protein^14^. This region is followed by a coiled-coil domain of varying lengths: Short in TC3 and TC4, and elongated in TC9, which may include Laminin I and helix-rich Mycoplasma domains. All fibers in these classes have a β-helical C-terminal domain, some of which with homology to fibronectins. TC4 and TC9 often present a β-sandwich C-terminal head (pseudo-domain D69), that has structural homology with lectins and GOLD domains.

TC7 consists of three fibers (RBP_13, RBP_33, and RBP_60) from three different sequence classes. Although the mean TM score of this class is greater than 0.4, RBP_60 is an outlier (**Fig. 3b, Supplementary Fig. 9c**). RBP_60 features a gp37 needle domain but has a completely different C-terminal part compared to TC6. The other TC7 fibers share a similar global architecture, with a coiled-coil N-terminal region and a C-terminal peptidase S74 domain. However, they share only 25.7% sequence identity.

TC8 is found in *Vequintavirinae* phages and was present in both Bas49 and Bas54 (**Fig. 3c**). These fibers can be divided into three pseudo-domains: an N-terminal α-helical bundle (D15), two consecutive β-helices (D32) and a C-terminal head (D31). The region D15 could not be fitted in the cryo-EM map, which could mean that it adopts a different conformation when attached to the virion, or it is cleaved during virion assembly. D31 seemed to be connected to another density, for which no fiber could be fitted. This indicates that D31 can be an adaptor for a tail tip adhesin, or for a longer tail fiber complex as in T4 LTF. (**Supplementary Fig. 4**).

TC12 consists of several subdomains that appear to be permutable, insertable, or replaceable (**Supplementary Fig. 9d**). RBP_10 shares the same domain order as RBP_11 and RBP_57, except for a single replacement at the C-terminus. This replacement was confirmed by cryo-EM (**Fig. 3d)**. RBP_11 and 57 have a peptidase S74 domain (D21) at their C-terminus, whereas RBP_10 has a lectin-like head (D69). This can indicate that *Escherichia phage* MaxTheCat uses a gp38-like adhesin for binding whereas *Escherichia phage* EmilHeitz uses the D69 region. As expected, the cryo-EM map of RBP_11 neither shows any density for the peptidase domain, nor any density for an adhesin protein (**Supplementary Fig. 4**). RBP_49 is the only TC12 member from a different sequence class, and it has a distinct N-terminal region, an elongated coiled-coil with a carboxypeptidase hit at its ends, that shares structural homology with *Vibrio* phage XM1 collar protein (7KJK). This indicates that RBP_49 can be located towards the neck region of its phage. RBP_49 lacks one copy of D42, the D7 pseudo-domains, and a non-classified region found between D7 and D42 in the other members of TC12. On Bas49 and Bas54, these pseudo-domains are in the interaction region with the other baseplate fibers (**Supplementary Fig 4**.), thus their absence in RBP_49 might be related to the location of the fiber.

TC16 has three fibers with elongated coiled-coil regions. Two fibers (RBP_52 and RBP_50) share a similar T7 N-terminal adaptor (D66), and a β-helical terminus followed by different types of receptor heads (T7-like, and Mu-like). The β-helical pseudo-domain D25 found in RBP_50 is a well spread motif also found on TC12, TC13 and TC14.

Most D-classes preserved their relative positions in different RBPs, but we identified five mobile pseudo-domains (D7, D11, D40, D42, and D73; **Supplementary Fig. 10a, b**). Four of these were previously classified: D7 is a "Pyocin knob," and D11, D40 and D73 are present in the distal region of T4’s gp34. However, D42 represents a novel domain. We confirmed the existence of D42 in Bas54 and Bas49 (**Fig. 4d**). HMM profiles of D7 and D42 retrieved tail fibers belonging to six and four different TCs, respectively, and they can be found in diverse phage subfamilies (**Supplementary Fig. 10c,d**). While the function of these highly abundant domains remains to be determined, their location within modular fibers—particularly in fibers with varying N-terminal regions, such as in different TC12 members—suggests that the gene encoding these domains may facilitate the lateral transfer of N-terminal domains.

**Fig. 4.**
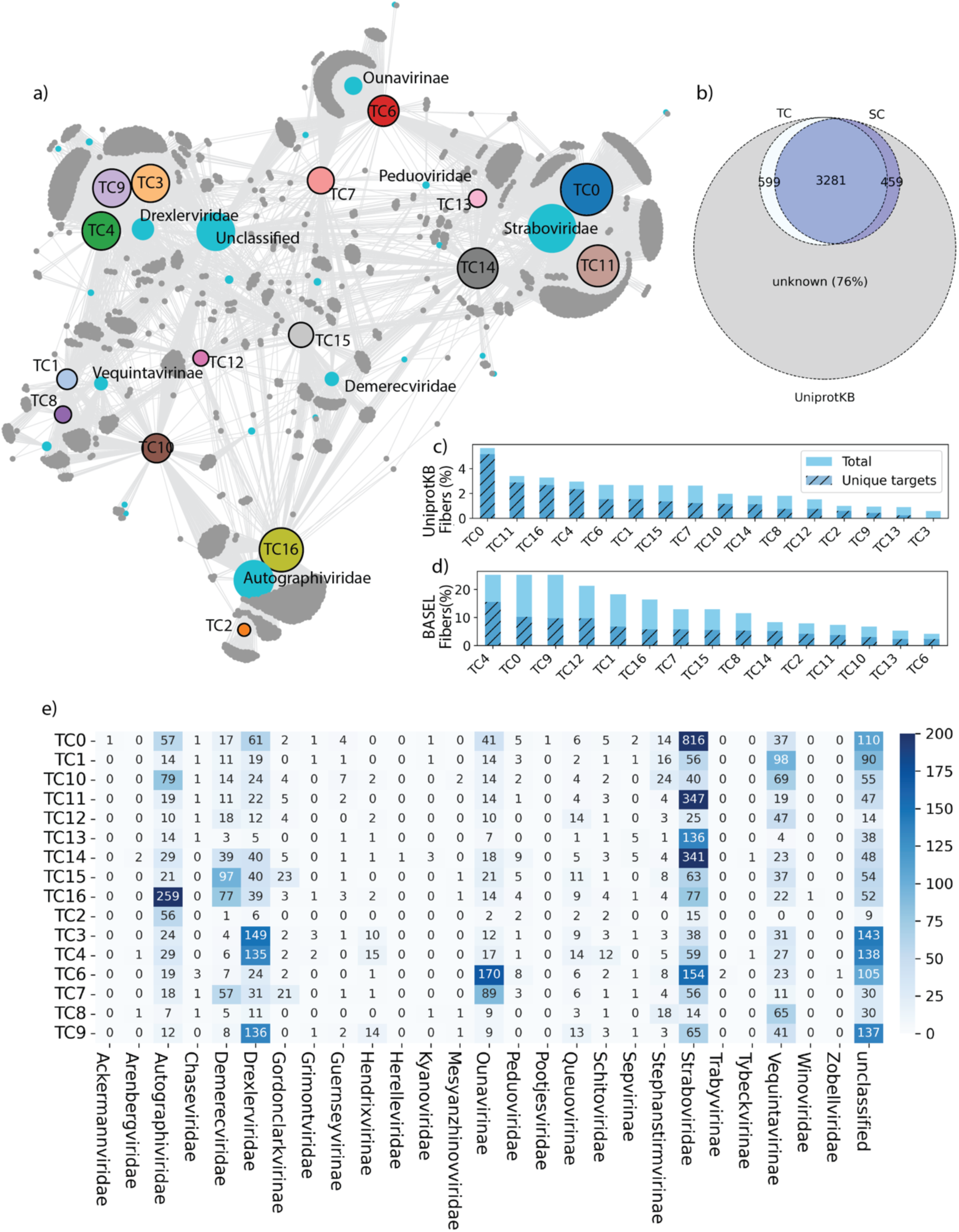
– The partial phage tail fiber atlas. a) Network connecting the TC with phage family or subfamily. Node size is proportional to number of proteins retrieve for each label. In turquoise: phage classification; in dark gray: phage name. b) Venn-diagram representing the total HMMsearch hits of all TC (light blue) and sequence classes (dark blue) against all annotated phage tail fibers in UniProtKB (gray). c,d) Histogram of total (skyblue) and unique (hashed/dark blue) targets retrieved for each TC against the BASEL collection tail fibers (c) and the UniProtKB tail fiber search (d). e) Heatmap of distribution of TC classes among different phage families and subfamilies.

### III.6 The new TC describes 24% of the tail fiber atlas

To verify the representability of our structural classes, we selected 16,345 annotated fiber sequences from UniProtKB. An initial query for ‘tail fiber’ yielded over 64,000 results. We filtered these by selecting only *Caudoviricetes* proteins with lengths between 300 and 3,000 residues and excluded those with ‘chaperone’ in their names. We then created HMM profiles from our 16 TC classes and ran these against the selected fibers (**Fig. 4**). The same procedure was applied to the sequence classes.

From the structural classes, we found a total of 6,739 high-confident matches (E-value < 10^-10^), corresponding to 4,417 unique proteins, which represents 24.0% of the total targets (**Fig. 4b,c**). Using the sequence clusters, we recovered 23.0% of the total targets. This partial coverage was expected, as our dataset primarily consisted of *E. coli* phages and our structural classes did not include common RBPs such as depolymerases. Moreover, there can still be sequence-diverse fibers with conserved structure that were not detected by the HMM profiles. Similar analysis against the RBP set from the BASEL collection retrieved 95% of the sequences. This value was expected to be lower than 100%, as we excluded the TC5 and TC17 from the search (**Fig. 4b**).

Notably, most retrived sequences from UniprotKB belonged to TC0 fibers (**Fig. 4d-e**), and the *Straboviridae* subfamily. This apparent abundance could also be due to a natural unbalance in the dataset towards the most well-studied phages, such as T4. A great number of fiber hits for all classes belonged to phages that are as of yet unclassified.

## IV. Conclusions

This study presents a generalized method for predicting tail fiber structures, and a novel way of classifying these predictions. This approach allows us to not only explore the diversity of the known tail fiber universe but can also be used to hint at possible evolutionary paths taken by those viruses. We established a robust pipeline to model, classify, and analyze the structure of those protein complexes. This pipeline was validated on three well-studied phages by using single-particle cryo-EM. We found 15 structural classes representing 24% of the known tail fiber universe. Our methodology and findings set the stage for the development of a complete tail fiber atlas, offering valuable insights into bacteriophage diversity and evolution. Furthermore, it might suggest strategies for tail fiber modification in phage applications in biotechnology and biomedicine.

## Data and Code Availability

The RBPseg pipeline, the scripts used to analyze the data and the RBPseg/AF2M models present in this manuscript are public available at http://github.com/vkleinsousa/RBPseg.

The Cryo-EM maps were deposited at EMDB. Entries: EMD-51870 (Bas36), EMD-51868 (Bas49), EMD-51869 (Bas54).

## Acknowledgments

The Novo Nordisk Foundation Center for Protein Research is supported financially by the Novo Nordisk Foundation (NNF14CC0001). N.M.I.T. acknowledges support from an NNF Hallas-Møller Emerging Investigator grant (NNF17OC0031006), an NNF Hallas-Møller Ascending Investigator grant (NNF23OC0081528) and an NNF Project grant (NNF21OC0071948) and is also a member of the Integrative Structural Biology Cluster (ISBUC) at the University of Copenhagen. V.K-S acknowledges the Novo Nordisk Foundation Copenhagen PhD Programme for grant NNF0069780. We acknowledge the Core Facility of Integrated Microscopy at University of Copenhagen (CFIM) for help with data collection. We acknowledge the Big Data Management Platform at Novo Nordisk Foundation Center for Protein Research for the computational resources.

## Author contribution

V.K-S., N.M.I.T and C.S.K. conceived the project. V.K-S. developed the computational framework, models, bioinformatical and statistical analysis. V.K-S. conceptualized the sDp and SC methods. V.K-S. and A.R-E. purified the bacteriophages. V.K-S., A.R-E. and N.S. prepared Cryo-EM sample and collected data. V.K-S. and A.R-E. processed cryo-EM datasets. V.K-S and C.S.K. cross-validated models and maps. V.K-S. wrote the manuscript and prepared figures, with input from all the authors. All the authors contributed for the revision of the manuscript.

## Use of large language models

Large language models (ChatGPT and Copilot) were used to enhance text readability and code debugging/annotation.

## Supplementary Materials

**Supplementary Figure 1.** RBPseg workflow in detail, step-by-step demonstrating the architecture of RBPseg using TC14 fiber as example. A FASTA file is input to ESMfold, which generates a monomeric model. This model is fractioned in the sDp module. Fraction FASTA files are modeled using AF2M, merged and relaxed.

**Supplementary Figure 2.** Selection of tail fiber representatives. a) Plot showing the consensus between PhANNs and PhageRBPdetect (orange). PhANNS fibers were selected as positive tail fiber when metric was greater than 8. PhageRBPdetect positive fibers, but PhANNs negative fibers are shown in yellow. b) Sequence identity matrix of all double positive fibers. c-d) Hierarchical clustering of tail fibers based on sequence identity shows for major groups. e) Sequence identity matrix of 67 selected fibers.

**Supplementary Figure 3.** Generality of sDp and its variants. **(a)** Comparison of PAE and sDp variants for the largest protein in the AlphaFold *E. coli* database (AF-P76347). The first row displays heatmaps of the PAE (left) and sDp all-against-all matrices (right). The second row shows a 2D UMAP projection of these matrices, colored by domain clusters identified using HDBSCAN with a minimum cluster distance of 40. **(b)** Structure of AF-P76347, colored according to domain organization. **(c)** Plot showing the absolute correlation between sDp variants and PAE for AF-P76347 across different Pair Distance Constants (D). (**d)** Pearson correlation between sDp variants and PAE for 1,000 randomly selected representatives from the Alphafold *E. coli* database, plotted against different values of D. A 4th-degree polynomial fit indicates a maximum (highlighted in yellow) at D = 9.87, with a correlation of −0.71. The fit yields R-squared = 0.9988, RMSE = 0.0047, and MAE = 0.0040. **(e-f)** Distribution of individual correlations depending on sequence length and mean PAE at D = 10. The p-values represent the results of a one-tailed Mann-Whitney U test. **(g)** Set of random representatives from the dataset. The first row shows sDp matrices, the second row shows PAE matrices, the third row presents sDp x PAE correlations, the fourth row depicts AF models colored by domain organization, and the fifth row displays UMAP projections of sDp matrices colored by protein regions.

**Supplementary Figure 4.** Map-model cross-correlation for all experimented validated TC versus residue number, and Bas54 refined maps. (a) The cross-correlation values plotted against the residue number for all experimentally validated TCs. The red line indicates the threshold at y = 0.5. The cross-correlation of the map against the full RBPseg model is shown in ‘sky blue’ (dash-dot line), while the cross-correlation against pseudo-domains, calculated using the sDp approach, is depicted in ‘green’ (dashed line). (b) Box plot showing the cross-correlation values for each comparison. *** denotes a p-value < 0.001 from a Z-test comparing full RBPseg models with individual pseudo-domains, while ‘ns’ indicates comparisons that are not statistically significant. (c) Refined Bas54 composite homogeneous and local refined mapes maps (EMD-51869, contour levels: 1.4 and 0.8) with superimposed pseudo-domains for RBP04, RBP11, and RBP65. Scale bar represents 10 nm. (d) Missing N-terminal region of RBP65, which is present in all members of TC8 (EMD-51869, contour levels: 2.0). (e) C-terminal region of RBP11. In ‘green,’ the Pfam-annotated chaperone of endosialidase is highlighted, while in ‘sky blue,’ the coiled-coil C-terminus is shown. Both regions are absent in the mature phage (EMD-51869, contour levels: 0.2).

**Supplementary Figure 5.** The TC classes and its members (part 1) colored based on sequence conservation. The sequence conservation of each TC class and its members is depicted, with coloring based on the degree of conservation. Sequence conservation was calculated using the multiple sequence alignment (MSA) of each TC class, highlighting regions of high (1) and low (−3) conservation across the members.

**Supplementary Figure 6.** The TC classes and its members (part 2). Analysis of classes TC5, TC17, TC7, TC12. (a) and (b) TC5 and TC17 exhibited low interclass mean TM scores and low sequence conservation, with proteins colored based on sequence conservation. (c) Members of TC7 are colored based on sequence conservation. RBP_13 and RBP_33 share common domains: D21 and D42 (with different copy numbers). RBP_60 shows no structural or sequence similarity with other class members and was likely misclassified; it shares a needle domain with TC6. (d) A detailed analysis of the modular class TC12, with proteins colored based on sequence conservation. Three members share the same N-terminal adaptor, while RBP_49 has a different N-terminal, lacks D7 domains, and contains one less copy of D42.

**Supplementary Figure 7.** Clustering criteria for TCs. a) TM matrix all-against-all, color scale represents the mean TM score between both fibers. b) Inner cluster mean TM score (TM, blue), standard deviation of icTM (sTM, green), maximum size of the clusters divided by 100 (yellow), Silhouette score (red) against the cluster number (#clusters). The dots represent the average of 10 independent runs for each cluster number. c) Structural clustering similarity metric (SM blue dots), and predicted exponential decay fit (pSM, red) against #clusters. d) Double derivative of SM and pSM (*pSM*′′). Green line represents number of clusters = 18 (*pSM*^))^ < 0.001).

**Supplementary Figure 8.** Clustering criteria for pseudo-domains and classes metrics. (a) Distribution of pseudo-domain sizes after applying the sDp approach with spectral clustering, using an expected domain size of 120 residues. Segments smaller than 20 residues or larger than 400 residues were excluded from the classification. (b) SM and pSM scores for the pseudo-domain classification. (c) Left: The mean TM-score for each pseudo-domain class (D classes), with reference lines indicating important thresholds—‘sky blue’ dashed line represents the random class mean TM-score, ‘red’ dashed line marks a TM-score of 0.4, and the ‘black’ dashed line represents the overall average mean TM-score. Center: The relative position of each pseudo-domain within the original RBP, calculated as the last residue number of each pseudo-domain divided by the C-terminal residue of the full-length RBP. Right: The length (in residues) of each pseudo-domain.

**Supplementary Figure 9.** Domain annotations of various RBPs across different TCs using InterPro (functional annotation) and Foldseek (PDB structure matches). The full-length RBP sequences are represented in ‘purple’ (solid line), indicating the entire protein. InterPro annotations were retrieved for the full-length RBP sequences and filtered with an e-value threshold of < 1e-5 to highlight functionally relevant domains. These functional domains are depicted along the RBP sequences with color-coded segments. Foldseek analysis identifies structural matches from PDB entries, with two levels of alignment confidence: high-confidence alignments (TM-score > 0.8) are shown in ‘dark blue’ (solid line), while moderate-confidence alignments (TM-score > 0.6) are indicated in ‘light blue’ (dashed line). The sDp-estimated pseudo-domains, which provide an approximation of possible domain boundaries within the RBPs, are shown in ‘sky blue’ (solid line).

**Supplementary Figure 10.** Mobile Pseudo-Domains. (a) Mean relative positions of all pseudo-domains with a mean TM-score > 0.6. Pseudo-domains found in different TC classes are highlighted in **green**. The mean relative position is calculated as the average of the relative position ((first_residue + last_residue) / (2 * RBP_length)) across all members of the same D-class. Error bars represent the standard deviation of the sample. (b) Overlap of all models belonging to the same D-class with a standard deviation in the mean position > 0.1. Annotations are based on sequence or structural matches (InterPro or Foldseek). The tail fiber atlas focuses on the analysis of mobile pseudo-domains (D7 in panel (c) and D42 in panel (d)), which are found in at least 4 TC classes.

**Supplementary Table 1. RBPseg and AF2M comparisons.**

**Supplementary Table 2. Cryo-EM data collection**

**Supplementary Table 3. TC classes.**

**Supplementary Table 4. D-classes and domain annotations.**

## VI. References

1. Klumpp, J., Dunne, M. & Loessner, M. J. A perfect fit: Bacteriophage receptor-binding proteins for diagnostic and therapeutic applications. Curr. Opin. Microbiol. 71, 102240 (2023).

2. Ouyang, R., Ongenae, V., Muok, A., Claessen, D. & Briegel, A. Phage fibers and spikes: a nanoscale Swiss army knife for host infection. Curr. Opin. Microbiol. 77, 102429 (2024).

3. Chen, P. et al. LamB, OmpC, and the Core Lipopolysaccharide of Escherichia coli K-12 Function as Receptors of Bacteriophage Bp7. J. Virol. 94, 10.1128/jvi.00325-20 (2020).

4. Taslem Mourosi, J., et al. Understanding Bacteriophage Tail Fiber Interaction with Host Surface Receptor: The Key “Blueprint” for Reprogramming Phage Host Range. Int. J. Mol. Sci. 23, 12146 (2022).

5. Dunne, M., Hupfeld, M., Klumpp, J. & Loessner, M. J. Molecular Basis of Bacterial Host Interactions by Gram-Positive Targeting Bacteriophages. Viruses 10, 397 (2018).

6. Greenfield, J. et al. Structure and function of bacteriophage CBA120 ORF211 (TSP2), the determinant of phage specificity towards E. coli O157:H7. Sci. Rep. 10, 15402 (2020).

7. Plattner, M. et al. Structure and Function of the Branched Receptor-Binding Complex of Bacteriophage CBA120. J. Mol. Biol. 431, 3718–3739 (2019).

8. Barbirz, S. et al. Crystal structure of Escherichia coli phage HK620 tailspike: podoviral tailspike endoglycosidase modules are evolutionarily related. Mol. Microbiol. 69, 303–316 (2008).

9. Pas, C., Latka, A., Fieseler, L. & Briers, Y. Phage tailspike modularity and horizontal gene transfer reveals specificity towards E. coli O-antigen serogroups. Virol. J. 20, 174 (2023).

10. Smug, B. J., Szczepaniak, K., Rocha, E. P. C., Dunin-Horkawicz, S. & Mostowy, R. J. Ongoing shuffling of protein fragments diversifies core viral functions linked to interactions with bacterial hosts. Nat. Commun. 14, 7460 (2023).

11. Haggård-Ljungquist, E., Halling, C. & Calendar, R. DNA sequences of the tail fiber genes of bacteriophage P2: evidence for horizontal transfer of tail fiber genes among unrelated bacteriophages. J. Bacteriol. 174, 1462–1477 (1992).

12. Taylor, N. M. I. et al. Structure of the T4 baseplate and its function in triggering sheath contraction. Nature 533, 346–352 (2016).

13. Chen, W., et al. Structural changes in bacteriophage T7 upon receptor-induced genome ejection. Proc. Natl. Acad. Sci. 118, e2102003118 (2021).

14. Linares, R. et al. Structural basis of bacteriophage T5 infection trigger and E. coli cell wall perforation. Sci. Adv. 9, eade9674 (2023).

15. Ge, X. & Wang, J. Structural mechanism of bacteriophage lambda tail’s interaction with the bacterial receptor. Nat. Commun. 15, 4185 (2024).

16. Hyman, P. & van Raaij, M. Bacteriophage T4 long tail fiber domains. Biophys. Rev. 10, 463–471 (2018).

17. Bartual, S. G. et al. Structure of the bacteriophage T4 long tail fiber receptor-binding tip. Proc. Natl. Acad. Sci. 107, 20287–20292 (2010).

18. Xiao, H. et al. Structure of the siphophage neck–Tail complex suggests that conserved tail tip proteins facilitate receptor binding and tail assembly. PLOS Biol. 21, e3002441 (2023).

19. Goulet, A., Spinelli, S., Mahony, J. & Cambillau, C. Conserved and Diverse Traits of Adhesion Devices from Siphoviridae Recognizing Proteinaceous or Saccharidic Receptors. Viruses 12, 512 (2020).

20. Šiborová, M. et al. Tail proteins of phage SU10 reorganize into the nozzle for genome delivery. Nat. Commun. 13, 5622 (2022).

21. Cunliffe, T. G., Parker, A. L. & Jaramillo, A. Pseudotyping Bacteriophage P2 Tail Fibers to Extend the Host Range for Biomedical Applications. ACS Synth. Biol. 11, 3207–3215 (2022).

22. Fa-arun, J., Huan, Y. W., Darmon, E. & Wang, B. Tail-Engineered Phage P2 Enables Delivery of Antimicrobials into Multiple Gut Pathogens. ACS Synth. Biol. 12, 596–607 (2023).

23. Altamirano, F. L. G. & Barr, J. J. Phage Therapy in the Postantibiotic Era. Clin. Microbiol. Rev. (2019) doi:10.1128/CMR.00066-18.

24. Jumper, J. et al. Highly accurate protein structure prediction with AlphaFold. Nature 596, 583–589 (2021).

25. Lin, Z. et al. Evolutionary-scale prediction of atomic-level protein structure with a language model. Science 379, 1123–1130 (2023).

26. Evans, R. et al. Protein complex prediction with AlphaFold-Multimer. 2021.10.04.463034 Preprint at 10.1101/2021.10.04.463034 (2022).

27. Bryant, P. et al. Predicting the structure of large protein complexes using AlphaFold and Monte Carlo tree search. Nat. Commun. 13, 6028 (2022).

28. Shor, B. & Schneidman-Duhovny, D. CombFold: predicting structures of large protein assemblies using a combinatorial assembly algorithm and AlphaFold2. Nat. Methods 21, 477–487 (2024).

29. Cambillau, C. & Goulet, A. Exploring Host-Binding Machineries of Mycobacteriophages with AlphaFold2. J. Virol. 0, e01793–22 (2023).

30. McInnes, L., Healy, J. & Melville, J. UMAP: Uniform Manifold Approximation and Projection for Dimension Reduction. Preprint at 10.48550/arXiv.1802.03426 (2020).

31. McInnes, L. & Healy, J. Accelerated Hierarchical Density Based Clustering. in 2017 IEEE International Conference on Data Mining Workshops (ICDMW) 33–42 (2017). doi:10.1109/ICDMW.2017.12.

32. Rousseeuw, P. J. Silhouettes: A graphical aid to the interpretation and validation of cluster analysis. J. Comput. Appl. Math. 20, 53–65 (1987).

33. Maffei, E. et al. Systematic exploration of Escherichia coli phage–host interactions with the BASEL phage collection. PLOS Biol. 19, e3001424 (2021).

34. The UniProt Consortium. UniProt: the Universal Protein Knowledgebase in 2023. Nucleic Acids Res. 51, D523–D531 (2023).

35. PhANNs, a fast and accurate tool and web server to classify phage structural proteins | PLOS Computational Biology. https://journals.plos.org/ploscompbiol/article?id=10.1371/journal.pcbi.1007845.

36. Boeckaerts, D., Stock, M., De Baets, B. & Briers, Y. Identification of Phage Receptor-Binding Protein Sequences with Hidden Markov Models and an Extreme Gradient Boosting Classifier. Viruses 14, 1329 (2022).

37. Sievers, F. & Higgins, D. G. Clustal Omega, Accurate Alignment of Very Large Numbers of Sequences. in Multiple Sequence Alignment Methods (ed. Russell, D. J.) 105– 116 (Humana Press, Totowa, NJ, 2014). doi:10.1007/978-1-62703-646-7_6.

38. Zimmermann, L. et al. A Completely Reimplemented MPI Bioinformatics Toolkit with a New HHpred Server at its Core. J. Mol. Biol. 430, 2237–2243 (2018).

39. Zhang, Y. & Skolnick, J. TM-align: a protein structure alignment algorithm based on the TM-score. Nucleic Acids Res. 33, 2302–2309 (2005).

40. Zhang, C., Shine, M., Pyle, A. M. & Zhang, Y. US-align: universal structure alignments of proteins, nucleic acids, and macromolecular complexes. Nat. Methods 19, 1109–1115 (2022).

41. ETE 3: Reconstruction, Analysis, and Visualization of Phylogenomic Data | Molecular Biology and Evolution | Oxford Academic. https://academic.oup.com/mbe/article/33/6/1635/2579822?login=true.

42. Steinegger, M. & Söding, J. MMseqs2 enables sensitive protein sequence searching for the analysis of massive data sets. Nat. Biotechnol. 35, 1026–1028 (2017).

43. Berman, H. M. et al. The Protein Data Bank. Nucleic Acids Res. 28, 235–242 (2000).

44. van Kempen, M. et al. Fast and accurate protein structure search with Foldseek. Nat. Biotechnol. 42, 243–246 (2024).

45. Paysan-Lafosse, T. et al. InterPro in 2022. Nucleic Acids Res. 51, D418–D427 (2023).

46. Wang, J. et al. The conserved domain database in 2023. Nucleic Acids Res. 51, D384– D388 (2023).

47. Marchler-Bauer, A. & Bryant, S. H. CD-Search: protein domain annotations on the fly. Nucleic Acids Res. 32, W327–W331 (2004).

48. ECOD: An Evolutionary Classification of Protein Domains | PLOS Computational Biology. https://journals.plos.org/ploscompbiol/article?id=10.1371/journal.pcbi.1003926.

49. Eddy, S. R. Accelerated Profile HMM Searches. PLOS Comput. Biol. 7, e1002195 (2011).

50. Plique, G. ipysigma. Zenodo 10.5281/zenodo.7521476 (2023).

51. Virtanen, P. et al. SciPy 1.0: fundamental algorithms for scientific computing in Python. Nat. Methods 17, 261–272 (2020).

52. Punjani, A., Rubinstein, J. L., Fleet, D. J. & Brubaker, M. A. cryoSPARC: algorithms for rapid unsupervised cryo-EM structure determination. Nat. Methods 14, 290–296 (2017).

53. Meng, E. C. et al. UCSF ChimeraX: Tools for structure building and analysis. Protein Sci. 32, e4792 (2023).

54. Afonine, P. V. et al. Real-space refinement in PHENIX for cryo-EM and crystallography. Acta Crystallogr. Sect. Struct. Biol. 74, 531–544 (2018).

55. Salomon-Ferrer, R., Case, D. A. & Walker, R. C. An overview of the Amber biomolecular simulation package. WIREs Comput. Mol. Sci. 3, 198–210 (2013).

56. Eastman, P. et al. OpenMM 7: Rapid development of high performance algorithms for molecular dynamics. PLOS Comput. Biol. 13, e1005659 (2017).

57. Tunyasuvunakool, K. et al. Highly accurate protein structure prediction for the human proteome. Nature 596, 590–596 (2021).

58. Schulz, E. C. et al. Crystal structure of an intramolecular chaperone mediating triple– β-helix folding. Nat. Struct. Mol. Biol. 17, 210–215 (2010).

59. Dunne, M., et al. *Salmonella* Phage S16 Tail Fiber Adhesin Features a Rare Polyglycine Rich Domain for Host Recognition. Structure 26, 1573–1582.e4 (2018).

60. Mori, Y. et al. Determination of the three-dimensional structure of bacteriophage Mu(-) tail fiber and its characterization. Virology 593, 110017 (2024).

61. North, O. I. Phage tail fibre assembly proteins employ a modular structure to drive the correct folding of diverse fibres. Nat. Microbiol. 4, 11 (2019).

62. Miller, J.-M., Knyazhanskaya, E. S., Buth, S. A., Prokhorov, N. S. & Leiman, P. G. Function of the bacteriophage P2 baseplate central spike Apex domain in the infection process. 2023.02.25.529910 Preprint at 10.1101/2023.02.25.529910 (2023).

