## Supplementary Figures and Supplementary Table 2 for "Towards a complete phage tail fiber structure atlas"

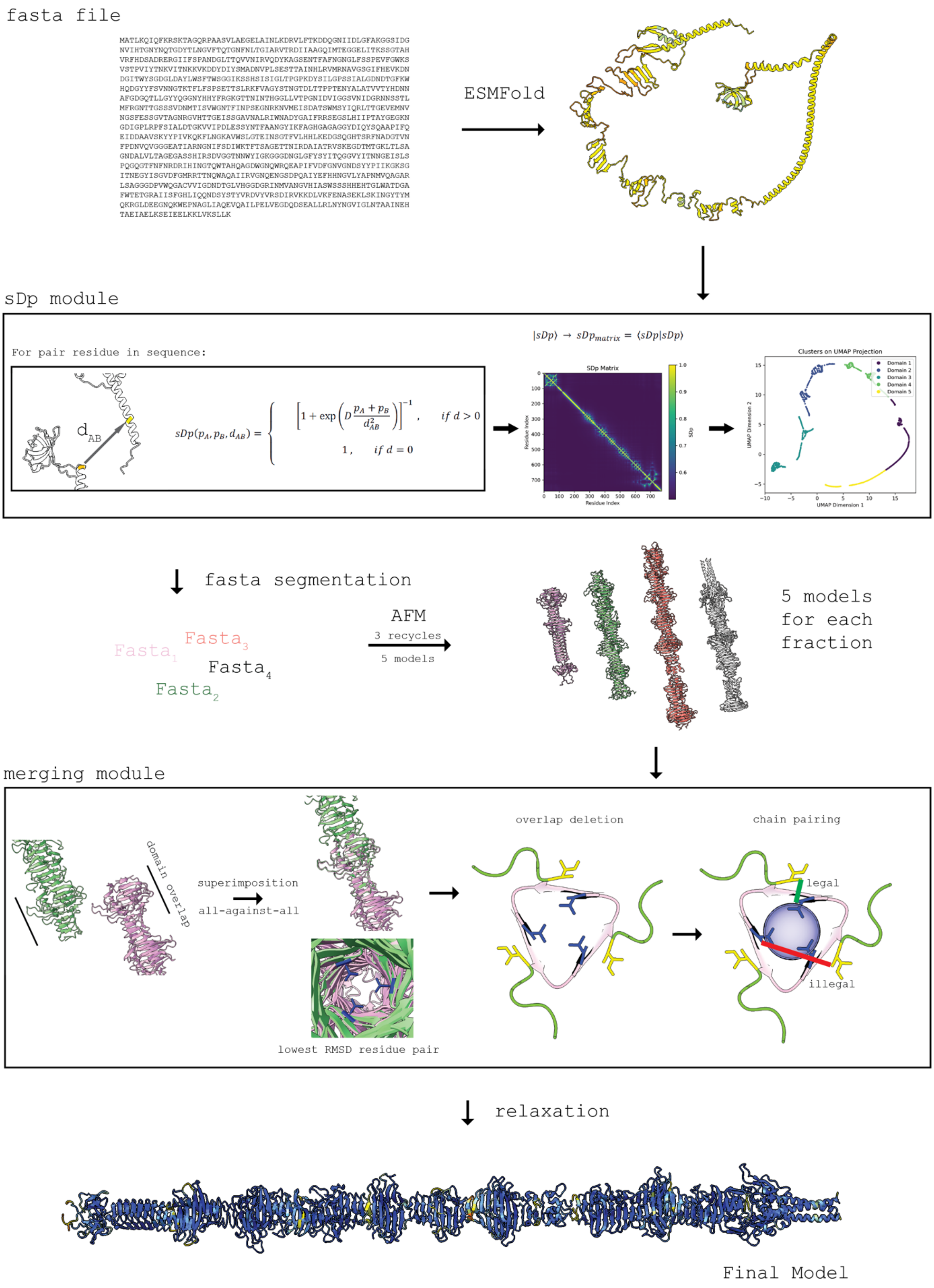


**Supplementary Figure 1.** RBPseg workflow in detail, step-by-step demonstrating the architecture of RBPseg using TC14 fiber as example. A FASTA file is input to ESMfold, which generates a monomeric model. This model is fractioned on the sDp module. Fraction FASTA files are modeled using AF2M, merged and relaxed.

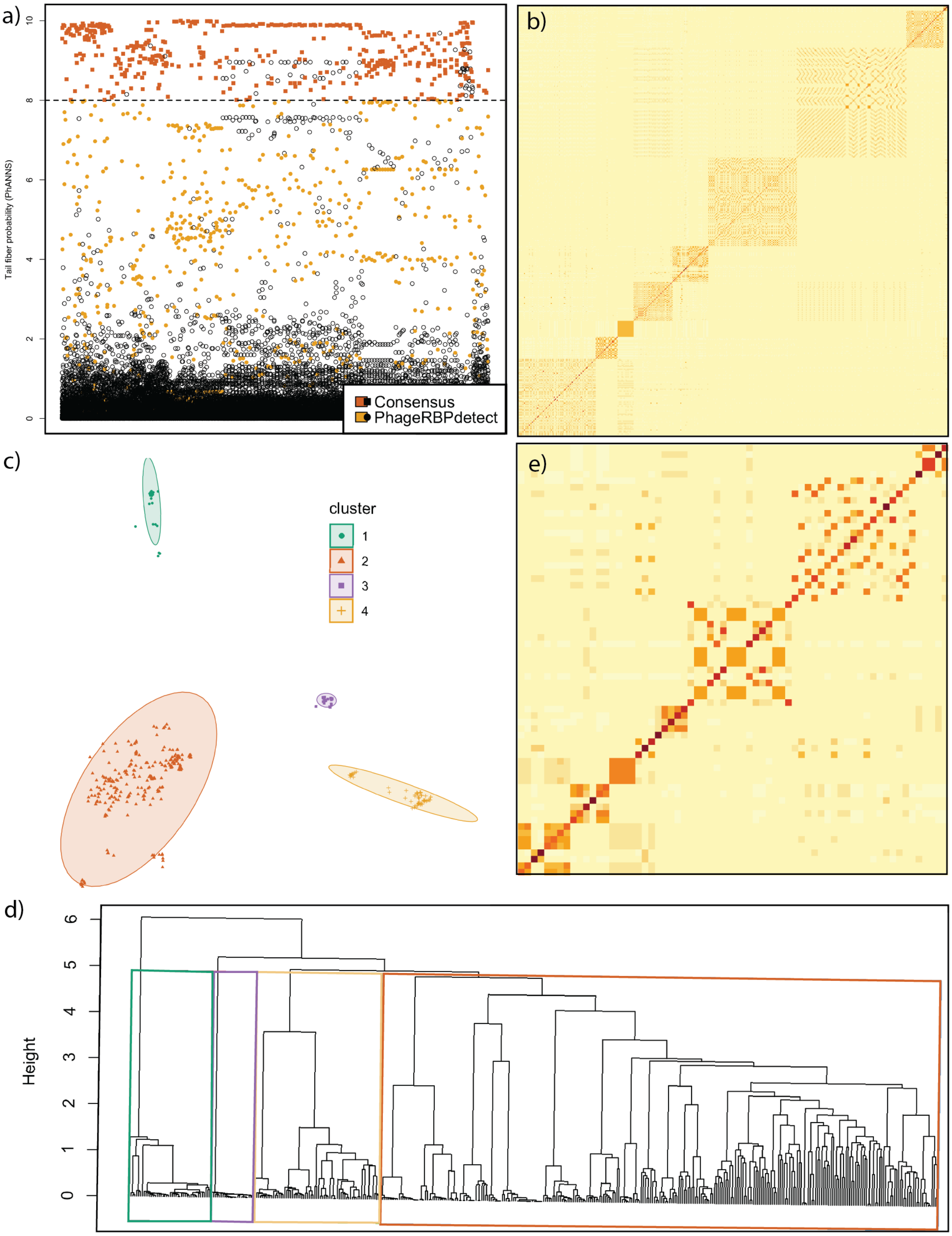


**Supplementary Figure 2.** Selection of tail fiber representatives. a) Plot showing the consensus between PhANNs and PhageRBPdetect (orange). PhANNS fibers were selected as positive tail fiber when metric was greater than 8. PhageRBPdetect positive fibers, but PhANNs negative fibers are shown in yellow. b) Sequence identity matrix of all double positive fibers. c-d) Hierarchical clustering of tail fibers based on sequence identity shows for major groups. e) Sequence identity matrix of 66 selected fibers.

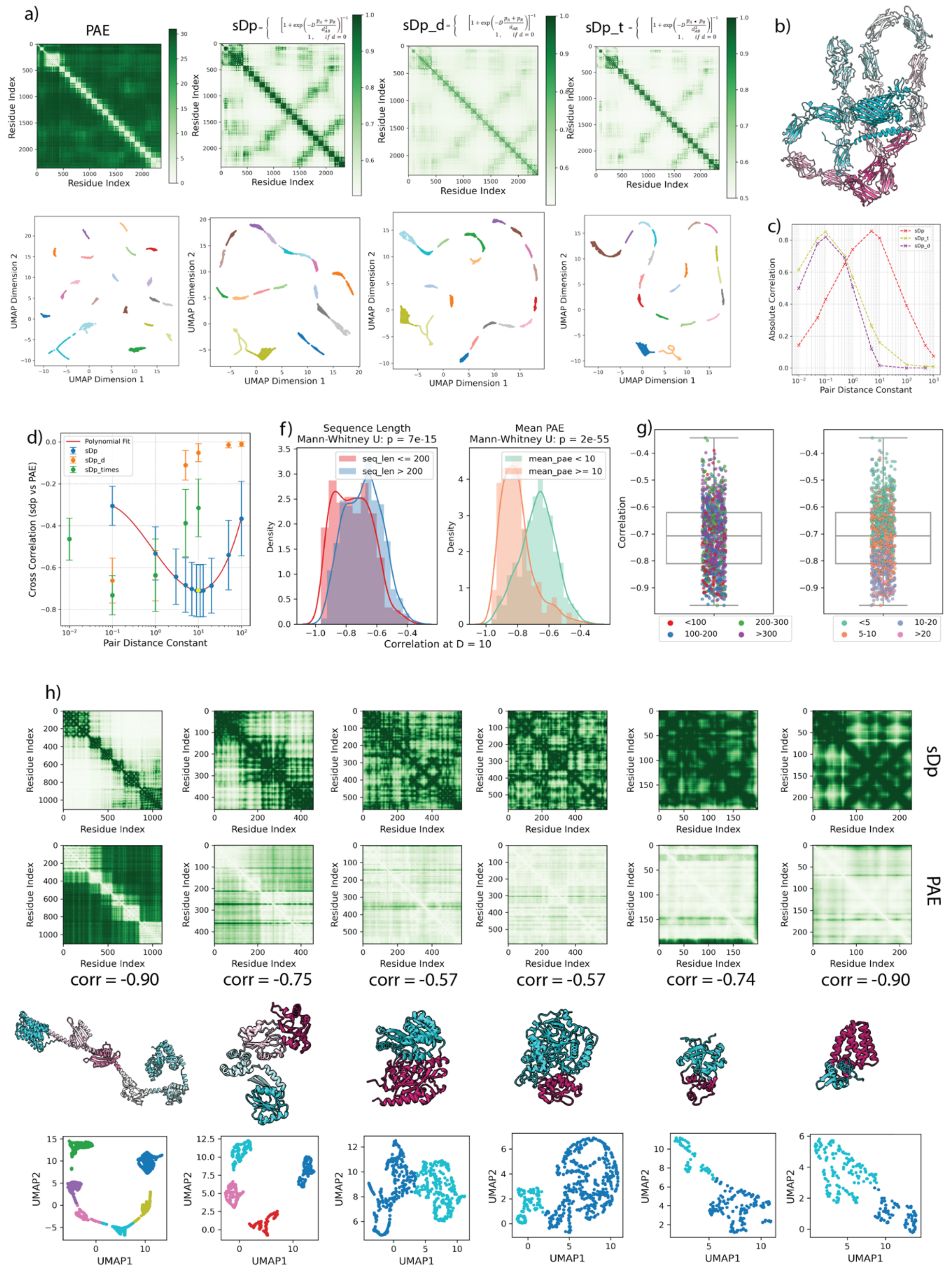


**Supplementary Figure 3.** Generality of sDp and its variants. **(a)** Comparison of PAE and sDp variants for the largest protein in the Alphafold *E. coli* database (AF-P76347). The first row displays heatmaps of the PAE (left) and sDp all-against-all matrices (right). The second row shows a 2D UMAP projection of these matrices, colored by domain clusters identified using

HDBSCAN with a minimum cluster distance of 40. **(b)** Structure of AF-P76347, colored according to domain organization. **(c)** Plot showing the absolute correlation between sDp variants and PAE for AF-P76347 across different Pair Distance Constants (D). (**d)** Pearson correlation between sDp variants and PAE for 1,000 randomly selected representatives from the Alphafold *E. coli* database, plotted against different values of D. A 4th-degree polynomial fit indicates a maximum (highlighted in yellow) at D = 9.87, with a correlation of -0.71. The fit yields R-squared = 0.9988, RMSE = 0.0047, and MAE = 0.0040. **(e-f)** Distribution of individual correlations depending on sequence length and mean PAE at D = 10. The p-values represent the results of a one-tailed Mann-Whitney U test. **(g)** Set of random representatives from the dataset. The first row shows sDp matrices, the second row shows PAE matrices, the third row presents sDp x PAE correlations, the fourth row depicts AF models colored by domain organization, and the fifth row displays UMAP projections of sDp matrices colored by protein regions.

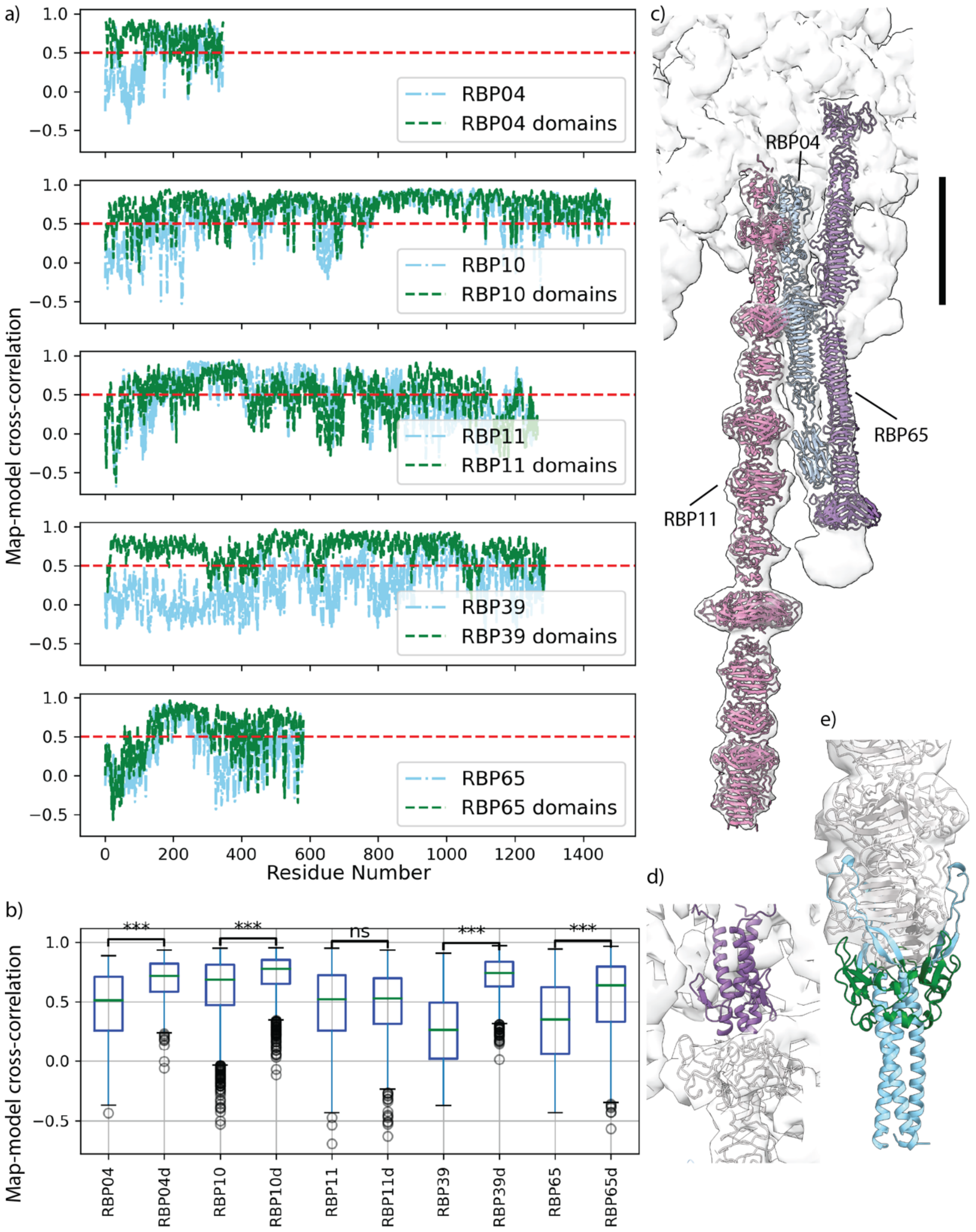


**Supplementary Figure 4.** Map-model cross-correlation for all experimented validated TC versus residue number, and Bas54 refined maps. (a) The cross-correlation values plotted against the residue number for all experimentally validated TCs. The red line indicates the threshold at y = 0.5. The cross-correlation of the map against the full RBPseg model is shown in 'sky blue' (dash-dot line), while the cross-correlation against pseudo-domains, calculated using the sDp approach, is depicted in 'green' (dashed line). (b) Box plot showing the cross-correlation values for each comparison. *** denotes a p-value < 0.001 from a Z-test comparing full RBPseg models with individual pseudo-domains, while 'ns' indicates comparisons that are not statistically significant. (c) Refined Bas54 composite homogeneous and local refined mapes maps (EMD-51869, contour levels: 1.4 and 0.8) with superimposed pseudo-domains for RBP04, RBP11, and RBP65. Scale bar represents 10 nm. (d) Missing N-terminal region of RBP65, which is present in all members of TC8 (EMD-51869, contour levels: 2.0). (e) C-terminal region of RBP11. In 'green,' the Pfam-annotated chaperone of endosialidase is highlighted, while in 'sky blue,' the coiled-coil C-terminus is shown. Both regions are absent in the mature phage (EMD-51869, contour levels: 0.2).

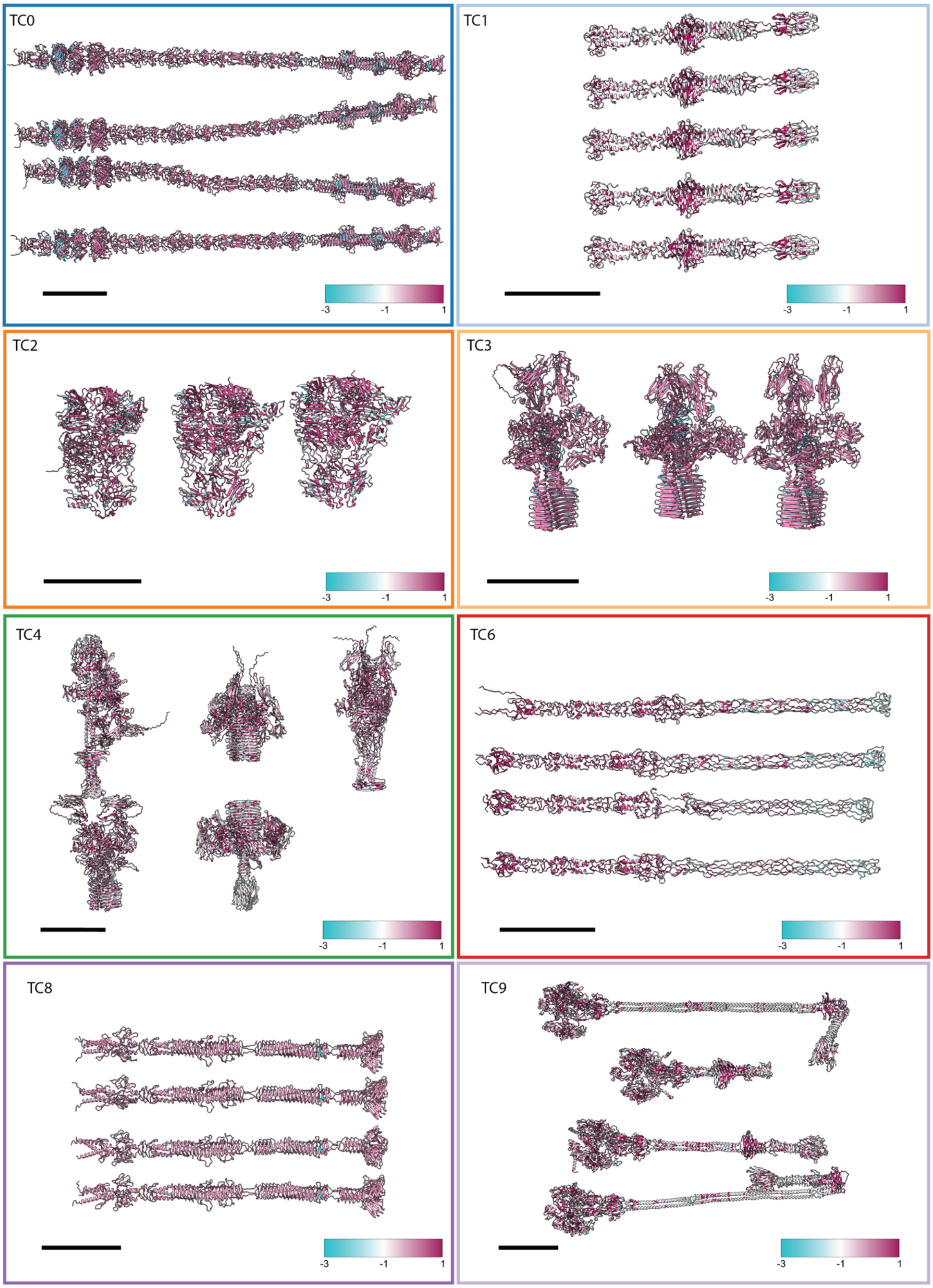
TC0
-3
-1
1
TC1
-3
-1
1
TC2
-3
-1
1
TC3
-3
-1
1
TC4
-3
-1
1
TC6
-3
-1
1
TC8
-3
-1
1
TC9
-3
-1
1

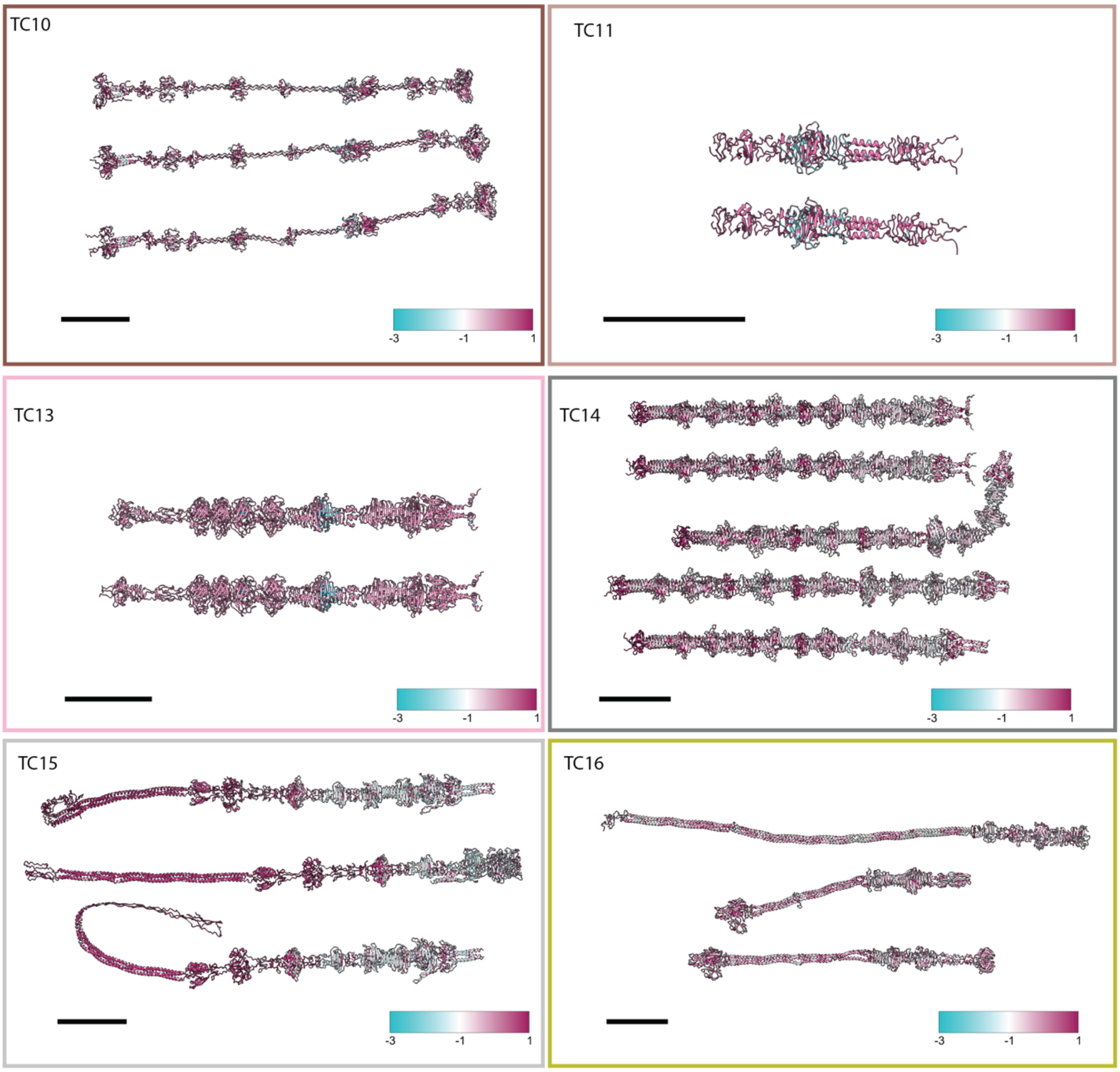


**Supplementary Figure 5.** The TC classes and its members (part 1) colored based on sequence conservation. The sequence conservation of each TC class and its members is depicted, with coloring based on the degree of conservation. Sequence conservation was calculated using the multiple sequence alignment (MSA) of each TC class, highlighting regions of high (1) and low (-3) conservation across the members.

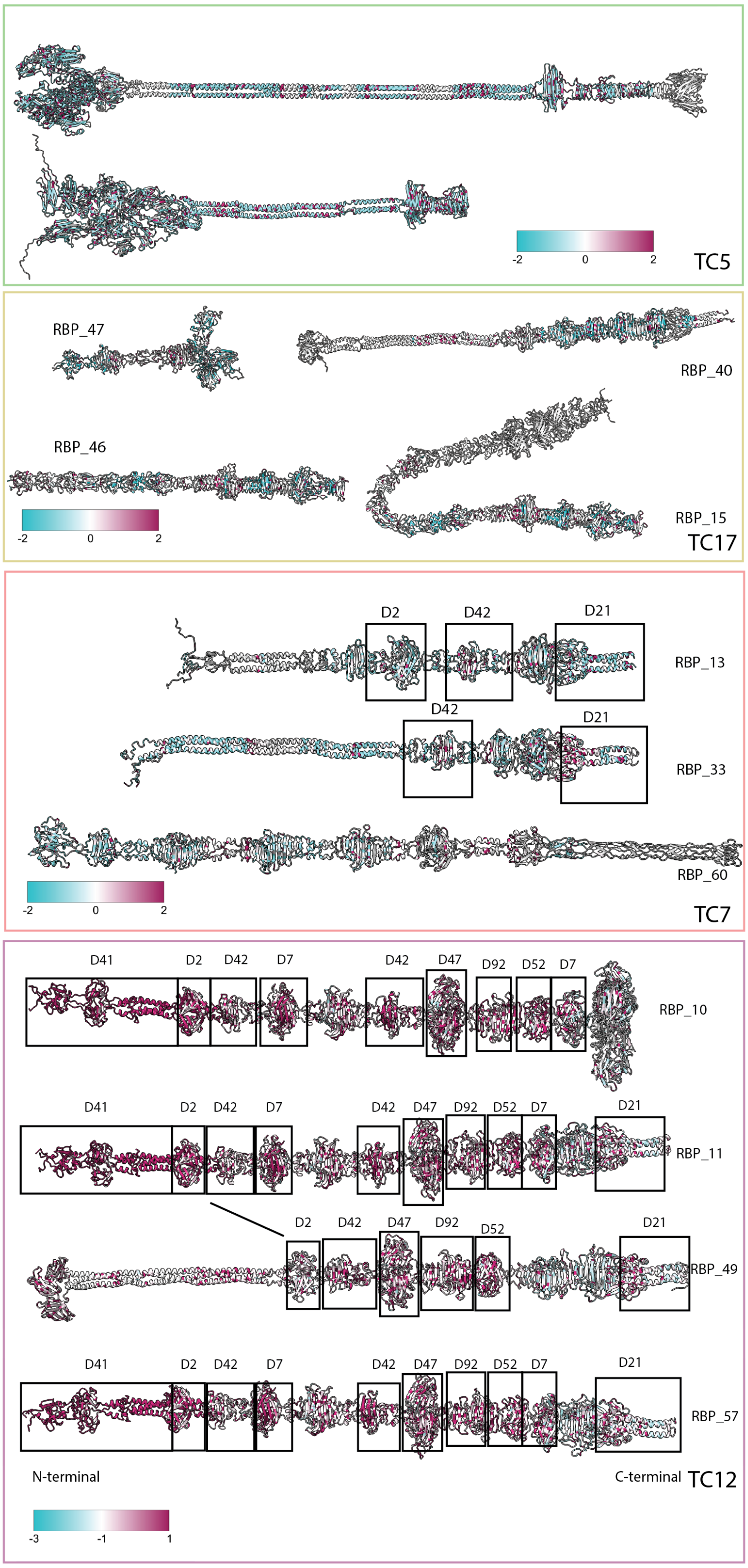


**Supplementary Figure 6.** Analysis of classes TC5, TC17, TC7, TC12. (a) and (b) TC5 and TC17 exhibited low interclass mean TM scores and low sequence conservation, with proteins colored based on sequence conservation. (c) Members of TC7 are colored based on sequence conservation. RBP_13 and RBP_33 share common domains: D21 and D42 (with different copy numbers). RBP_60 shows no structural or sequence similarity with other class members and was likely misclassified; it shares a needle domain with TC6. (d) A detailed analysis of the modular class TC12, with proteins colored based on sequence conservation. Three members share the same N-terminal adaptor, while RBP_49 has a different N-terminal, lacks D7 domains, and contains one less copy of D42.

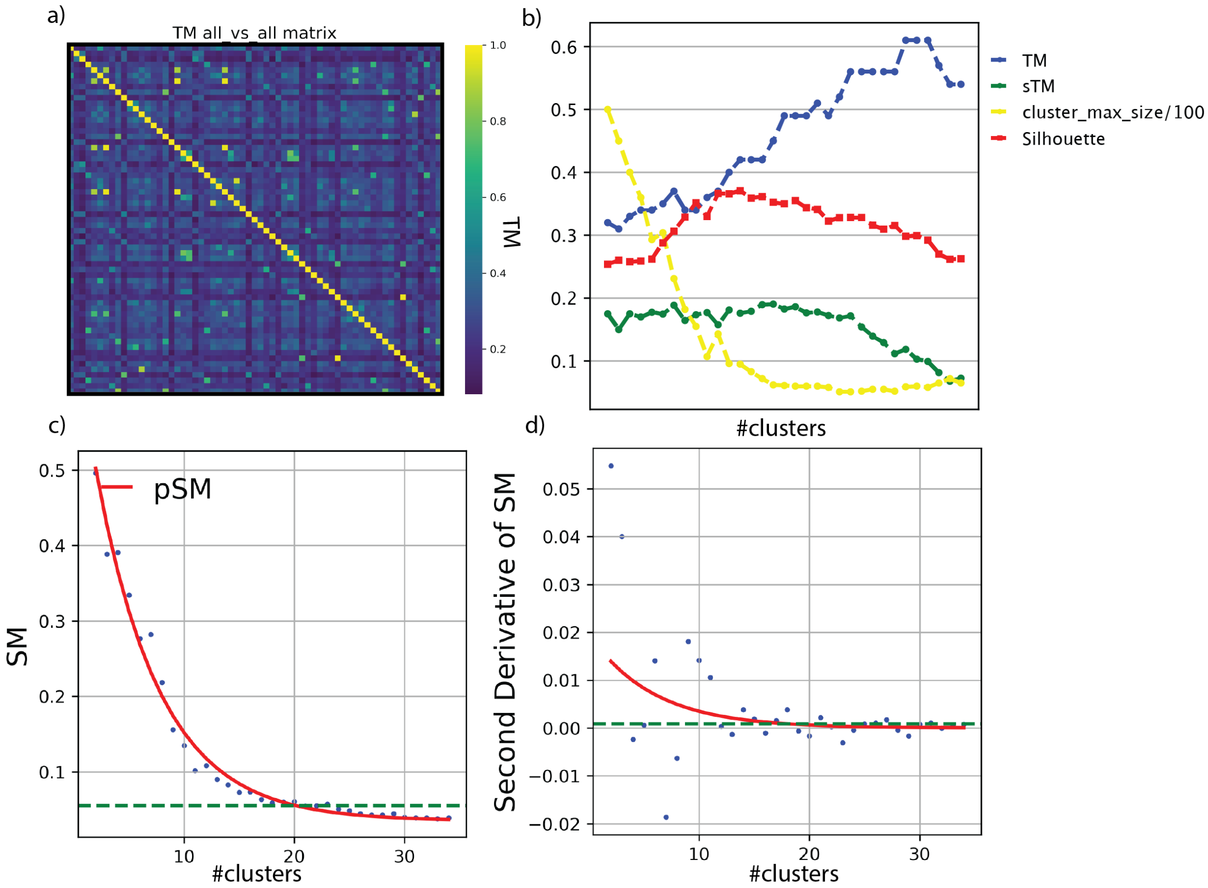


**Supplementary Figure 7.** Clustering criteria for TCs. a) TM matrix all-against-all, color scale represents the mean TM score between both fibers. b) Inner cluster mean TM score (TM, blue), standard deviation of icTM (sTM, green), maximum size of the clusters divided by 100 (yellow), Silhouette score (red) against the cluster number (#clusters). The dots represent the average of 10 independent runs for each cluster number. c) Structural clustering similarity metric (SM blue dots), and predicted exponential decay fit (pSM, red) against #clusters. d) Double derivative of SM and pSM ($pSM''$). Green line represents number of clusters = 18 ($pSM'' < 0.001$).

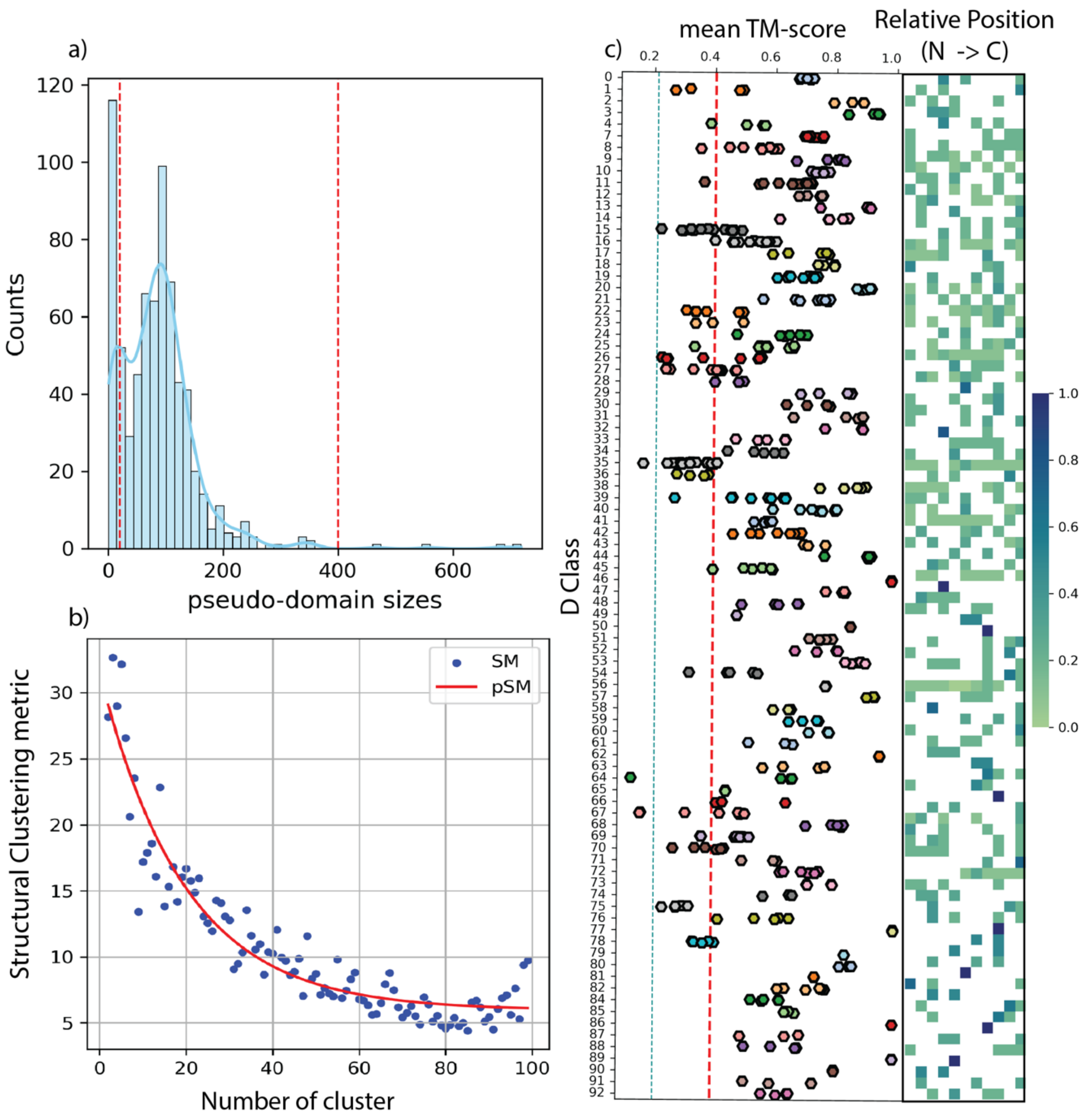


**Supplementary Figure 8.** Clustering criteria for pseudo-domains and classes metrics. (a) Distribution of pseudo-domain sizes after applying the sDp approach with spectral clustering, using an expected domain size of 120 residues. Segments smaller than 20 residues or larger than 400 residues were excluded from the classification. (b) SM and pSM scores for the pseudo-domain classification. (c) Left: The mean TM-score for each pseudo-domain class (D classes), with reference lines indicating important thresholds—'sky blue' dashed line represents the random class mean TM-score, 'red' dashed line marks a TM-score of 0.4, and the 'black' dashed line represents the overall average mean TM-score. Center: The relative position of each pseudo-domain within the original RBP, calculated as the last residue number of each pseudo-domain divided by the C-terminal residue of the full-length RBP. Right: The length (in residues) of each pseudo-domain.

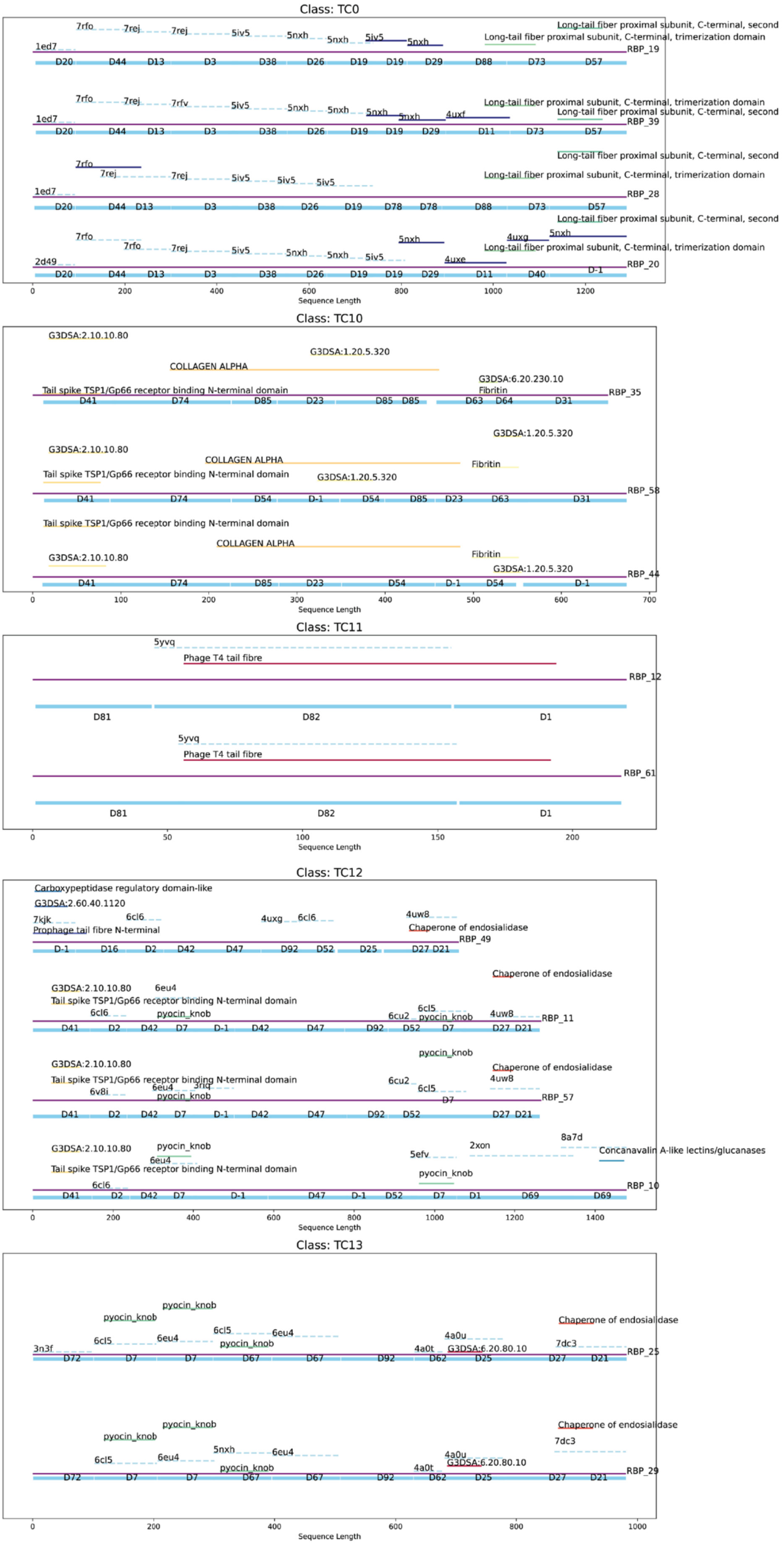

Class: TC0
7rfo
7rej
7rej
5iv5
5nxh
5nxh
5iv5
5nxh
Long-tail fiber proximal subunit, C-terminal, second
Long-tail fiber proximal subunit, C-terminal, trimerization domain
1ed7
RBP_19
D20 D44 D13 D3 D38 D26 D19 D19 D29 D88 D73 D57
7rfo
7rej
7rfv
5iv5
5nxh
5nxh
5nxh
5nxh
4uxf
RBP_39
D20 D44 D13 D3 D38 D26 D19 D19 D29 D11 D73 D57
7rfo
7rej
7rej
5iv5
5iv5
5iv5
RBP_28
D20 D44 D13 D3 D38 D26 D19 D78 D78 D88 D73 D57
5nxh
4uxg
5nxh
4uxe
2d49
RBP_20
D20 D44 D13 D3 D38 D26 D19 D19 D29 D11 D40 D-1
0 200 400 600 800 1000 1200
Sequence Length
Class: TC10
G3DSA:2.10.10.80
G3DSA:1.20.5.320
COLLAGEN ALPHA
G3DSA:6.20.230.10
Fibritin
Tail spike TSP1/Gp66 receptor binding N-terminal domain
RBP_35
D41 D74 D85 D23 D85 D85 D63 D64 D31
RBP_58
D41 D74 D54 D-1 D54 D85 D23 D63 D31
RBP_44
D41 D74 D85 D23 D54 D-1 D54 D-1
0 100 200 300 400 500 600 700
Sequence Length
Class: TC11
5yvq
Phage T4 tail fibre
RBP_12
D81 D82 D1
RBP_61
D81 D82 D1
0 50 100 150 200
Sequence Length
Class: TC12
Carboxypeptidase regulatory domain-like
G3DSA:2.60.40.1120
7kjk
6cl6
4uxg
6cl6
4uw8
Prophage tail fibre N-terminal
Chaperone of endosialidase
RBP_49
D-1 D16 D2 D42 D47 D92 D52 D25 D27 D21
G3DSA:2.10.10.80
6eu4
Tail spike TSP1/Gp66 receptor binding N-terminal domain
6cl6
pyocin_knob
6cl5
6cu2
4uw8
RBP_11
D41 D2 D42 D7 D-1 D42 D47 D92 D52 D7 D27 D21
6v8i
3rlq
6cl5
RBP_57
D41 D2 D42 D7 D-1 D42 D47 D92 D52 D27 D21
8a7d
2xon
5efv
Concanavalin A-like lectins/glucanases
RBP_10
D41 D2 D42 D7 D-1 D47 D-1 D52 D7 D1 D69 D69
0 200 400 600 800 1000 1200 1400
Sequence Length
Class: TC13
pyocin_knob
Chaperone of endosialidase
6cl5
6eu4
4a0u
7dc3
3n3f
4a0t
G3DSA:6.20.80.10
RBP_25
D72 D7 D7 D67 D67 D92 D62 D25 D27 D21
5nxh
RBP_29
D72 D7 D7 D67 D67 D92 D62 D25 D27 D21
0 200 400 600 800 1000
Sequence Length

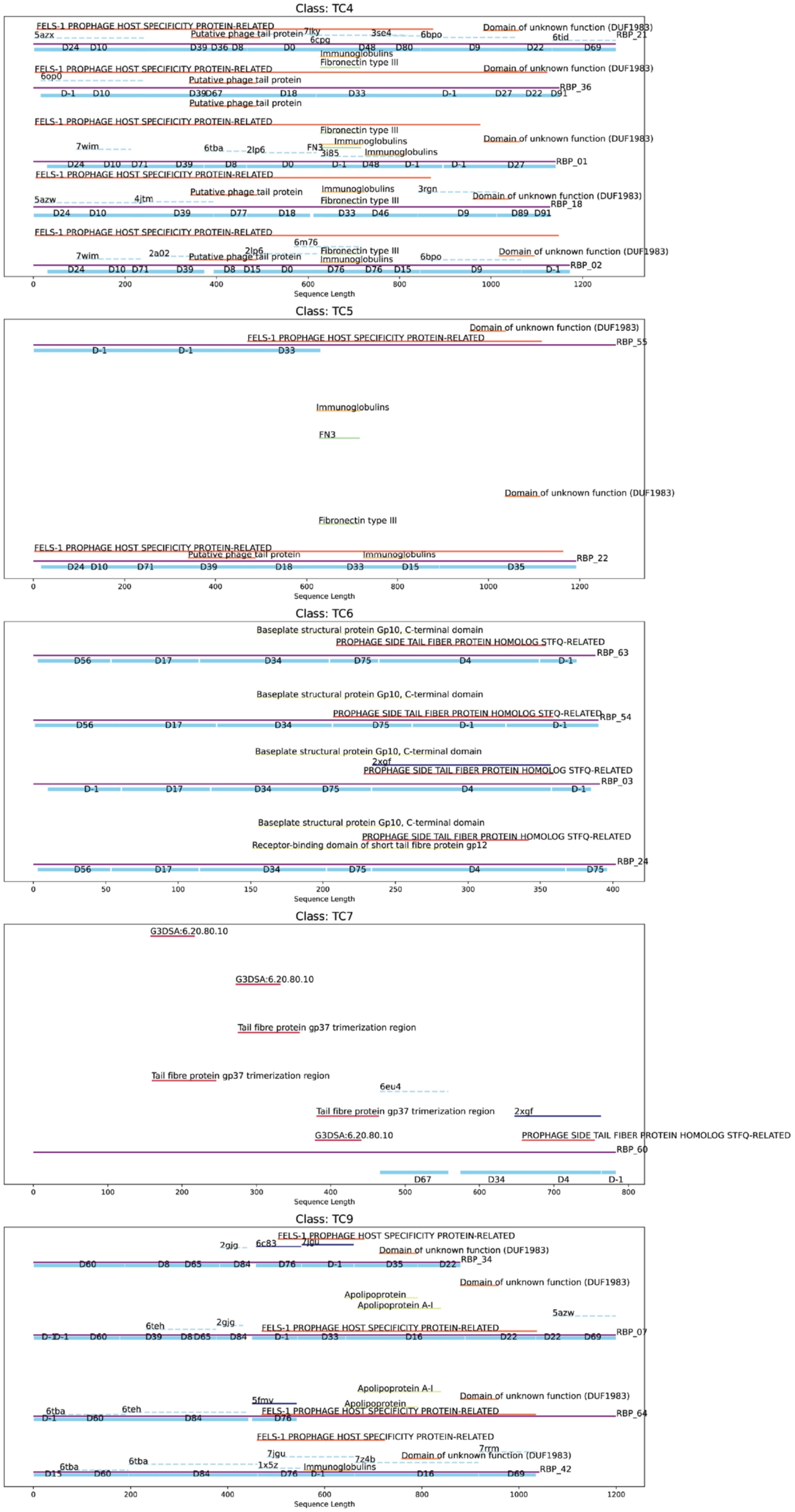
Class: TC4
FELS-1 PROPHAGE HOST SPECIFICITY PROTEIN-RELATED
Putative phage tail protein
Domain of unknown function (DUF1983)
Immunoglobulins
Fibronectin type III
FN3
RBP_21
RBP_36
RBP_01
RBP_18
RBP_02
Sequence Length
Class: TC5
Domain of unknown function (DUF1983)
FELS-1 PROPHAGE HOST SPECIFICITY PROTEIN-RELATED
RBP_55
Immunoglobulins
FN3
Fibronectin type III
Putative phage tail protein
RBP_22
Sequence Length
Class: TC6
Baseplate structural protein Gp10, C-terminal domain
PROPHAGE SIDE TAIL FIBER PROTEIN HOMOLOG STFQ-RELATED
RBP_63
RBP_54
RBP_03
Receptor-binding domain of short tail fibre protein gp12
RBP_24
Sequence Length
Class: TC7
G3DSA:6.20.80.10
Tail fibre protein gp37 trimerization region
PROPHAGE SIDE TAIL FIBER PROTEIN HOMOLOG STFQ-RELATED
RBP_60
Sequence Length
Class: TC9
FELS-1 PROPHAGE HOST SPECIFICITY PROTEIN-RELATED
Domain of unknown function (DUF1983)
Apolipoprotein
Apolipoprotein A-I
Immunoglobulins
RBP_34
RBP_07
RBP_64
RBP_42
Sequence Length

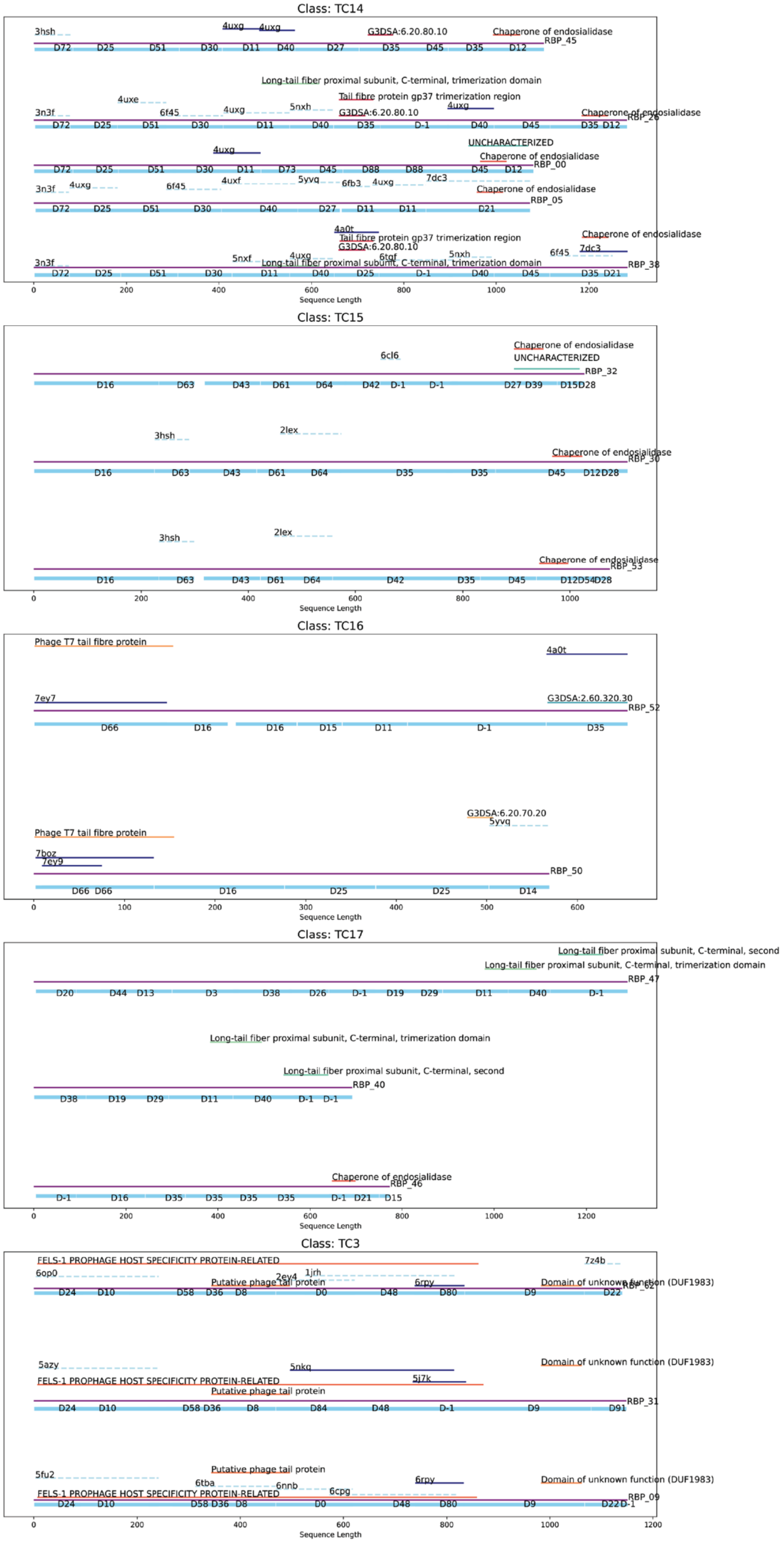
Class: TC14
3hsh
4uxg
4uxg
G3DSA:6.20.80.10
Chaperone of endosialidase
RBP_45
D72 D25 D51 D30 D11 D40 D27 D35 D45 D35 D12
Long-tail fiber proximal subunit, C-terminal, trimerization domain
4uxe
Tail fibre protein gp37 trimerization region
5nxh
4uxg
3n3f
6f45
4uxg
G3DSA:6.20.80.10
Chaperone of endosialidase
RBP_26
D72 D25 D51 D30 D11 D40 D35 D-1 D40 D45 D35 D12
4uxg
UNCHARACTERIZED
Chaperone of endosialidase
RBP_00
D72 D25 D51 D30 D11 D73 D45 D88 D88 D45 D12
3n3f
4uxg
6f45
4uxf
5yvq
6fb3
4uxg
7dc3
Chaperone of endosialidase
RBP_05
D72 D25 D51 D30 D40 D27 D11 D11 D21
4a0t
Tail fibre protein gp37 trimerization region
Chaperone of endosialidase
G3DSA:6.20.80.10
7dc3
5nxf
4uxg
6tgf
5nxh
6f45
3n3f
Long-tail fiber proximal subunit, C-terminal, trimerization domain
RBP_38
D72 D25 D51 D30 D11 D40 D25 D-1 D40 D45 D35 D21
0 200 400 600 800 1000 1200
Sequence Length
Class: TC15
6cl6
Chaperone of endosialidase
UNCHARACTERIZED
RBP_32
D16 D63 D43 D61 D64 D42 D-1 D-1 D27 D39 D15D28
3hsh
2lex
Chaperone of endosialidase
RBP_30
D16 D63 D43 D61 D64 D35 D35 D45 D12D28
3hsh
2lex
Chaperone of endosialidase
RBP_53
D16 D63 D43 D61 D64 D42 D35 D45 D12D54D28
0 200 400 600 800 1000
Sequence Length
Class: TC16
Phage T7 tail fibre protein
4a0t
7ey7
G3DSA:2.60.320.30
RBP_52
D66 D16 D16 D15 D11 D-1 D35
G3DSA:6.20.70.20
5yvq
Phage T7 tail fibre protein
7boz
7ey9
RBP_50
D66 D66 D16 D25 D25 D14
0 100 200 300 400 500 600
Sequence Length
Class: TC17
Long-tail fiber proximal subunit, C-terminal, second
Long-tail fiber proximal subunit, C-terminal, trimerization domain
RBP_47
D20 D44 D13 D3 D38 D26 D-1 D19 D29 D11 D40 D-1
Long-tail fiber proximal subunit, C-terminal, trimerization domain
Long-tail fiber proximal subunit, C-terminal, second
RBP_40
D38 D19 D29 D11 D40 D-1 D-1
Chaperone of endosialidase
RBP_46
D-1 D16 D35 D35 D35 D35 D-1 D21 D15
0 200 400 600 800 1000 1200
Sequence Length
Class: TC3
FELS-1 PROPHAGE HOST SPECIFICITY PROTEIN-RELATED
7z4b
6op0
1jrh
2ey4
Putative phage tail protein
6rpy
Domain of unknown function (DUF1983)
RBP_62
D24 D10 D58 D36 D8 D0 D48 D80 D9 D22
5azy
5nkq
Domain of unknown function (DUF1983)
FELS-1 PROPHAGE HOST SPECIFICITY PROTEIN-RELATED
5j7k
Putative phage tail protein
RBP_31
D24 D10 D58 D36 D8 D84 D48 D-1 D9 D91
5fu2
Putative phage tail protein
6rpy
Domain of unknown function (DUF1983)
6tba
6nnb
6cpg
FELS-1 PROPHAGE HOST SPECIFICITY PROTEIN-RELATED
RBP_09
D24 D10 D58 D36 D8 D0 D48 D80 D9 D22D-1
0 200 400 600 800 1000 1200
Sequence Length

**Supplementary Figure 9.** Domain annotations of various RBPs across different TCs using InterPro (functional annotation) and Foldseek (PDB structure matches). The full-length RBP sequences are represented in 'purple' (solid line), indicating the entire protein. InterPro annotations were retrieved for the full-length RBP sequences and filtered with an e-value threshold of < 1e-5 to highlight functionally relevant domains. These functional domains are depicted along the RBP sequences with color-coded segments. Foldseek analysis identifies structural matches from PDB entries, with two levels of alignment confidence: high-confidence alignments (TM-score > 0.8) are shown in 'dark blue' (solid line), while moderate-confidence alignments (TM-score > 0.6) are indicated in 'light blue' (dashed line). The sDp-estimated pseudo-domains, which provide an approximation of possible domain boundaries within the RBPs, are shown in 'sky blue' (solid line).

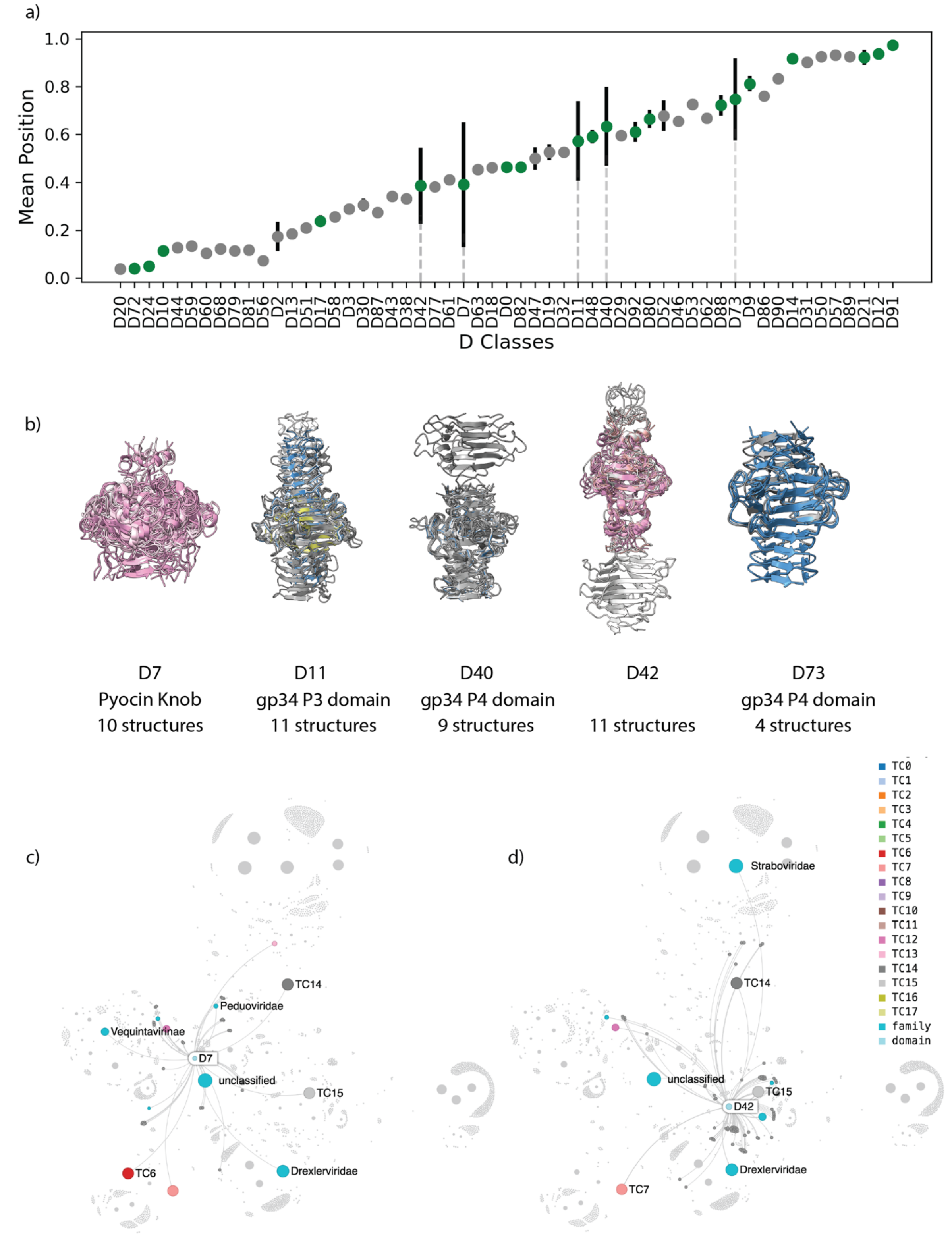


**Supplementary Figure 10. Mobile Pseudo-Domains.** (a) Mean relative positions of all pseudo-domains with a mean TM-score > 0.6. Pseudo-domains found in different TC classes are highlighted in **green**. The mean relative position is calculated as the average of the relative position ((first_residue + last_residue) / (2 * RBP_length)) across all members of the same D-class. Error bars represent the standard deviation of the sample. (b) Overlap of all models belonging to the same D-class with a standard deviation in the mean position > 0.1. Annotations are based on sequence or structural matches (InterPro or Foldseek). The tail fiber atlas focuses on the analysis of mobile pseudo-domains (D7 in panel (c) and D42 in panel (d)), which are found in at least 4 TC classes.

**Supplementary Table 2: Cryo-EM data collection**

| **Data collection and processing** | *Escherichia* phage Paracelsus (Bas36) | *Escherichia* phage MaxTheCat (Bas54) Baseplate | *Escherichia* phage MaxTheCat (Bas54) Distal Fiber | *Escherichia* phage EmilHeitz (Bas49) Baseplate | *Escherichia* phage EmilHeitz (Bas49) Distal Fiber |
|---|---|---|---|---|---|
| | ( EMD-51870) | (EMD-51869) | | (EMD-51868) | |
| Magnification Voltage (kV) | 300 | 300 | - | 300 | - |
| Electron exposure (e–/Å$^2$) | 40 | 40 | - | 40 | - |
| Defocus range (μm) | [-2,5;-0,5] | [-2,5;-0,5] | - | [-2,5;-0,5] | - |
| Pixel size (Å) – Data collection | 1,20 | 1,20 | - | 1,20 | - |
| Voxel size (Å) – Data processing | 4.6 | 4.6 | 4.6 | 3.6 | 3.6 |
| Symmetry imposed | C6 | C6 | C1 | C6 | C1 |
| Initial particle images (no.) | 8598 | 18179 | - | 16263 | - |
| Final particle images (no.) | 7986 | 17591 | - | 4833 | - |
| Map resolution (Å) (FSC threshold = 0.143) - mask | 9.46 | 9.46 | 9.46 | 7.32 | 7.67 |
| Map resolution range (no-mask, minimum until 25th percentile) (Å) | 10.55, 29.02 | 10.55, 27.82 | - | 8.23, 40.17 | - |
